# Distinct bacterial and protist plankton diversity dynamics uncovered through DNA-based monitoring in the Baltic Sea area

**DOI:** 10.1101/2024.08.14.607742

**Authors:** Krzysztof T Jurdzinski, Meike AC Latz, Anders Torstensson, Sonia Brugel, Mikael Hedblom, Yue O O Hu, Markus Lindh, Agneta Andersson, Bengt Karlson, Anders F Andersson

## Abstract

Planktonic microorganisms in coastal waters form the base of food web and biogeochemical cycles. The Baltic Sea area, with its pronounced environmental gradients, serves as a model coastal environment. Yet, microbial diversity assessment across these environmental gradients has so far lacked either taxonomic scope or the integration of spatial and temporal scales. Here, we analyzed protist and bacterial diversity using DNA metabarcoding across 398 samples synchronized with national monitoring of the Baltic Sea and the Kattegat-Skagerrak. We show that salinity, unlike other environmental factors, had a stronger effect on bacterial than on protist community composition. Likewise, Bayesian modeling showed that bacterial lineages were less likely than protists to occur in both lower (<9 PSU) and higher (>15 PSU) brackish salinities. Nonetheless, protist alpha diversity increased with salinity. Changes in bacterial alpha diversity were primarily seasonal and linked to influx of deepwater taxa through vertical mixing in winter. We propose that protists are ecologically less sensitive to salinity because compartmentalization allows them to disconnect basic metabolic processes from the cell membrane. Additionally, further and more frequent dispersal of bacteria might impede local adaptation. Ultimately, DNA-based environmental monitoring expands our understanding of microbial diversity patterns and the underlying factors.

## Introduction

Aquatic microorganisms play indispensable roles in biogeochemical cycles^1^. Phototrophic planktonic bacteria and protists account for around half of primary production on Earth^2^, with a disproportionately high contribution from coastal ecosystems^3^. More than a third of the human population lives near these ecosystems^4^ and depends on them^5,6^, while these environments experience extreme anthropogenic pressures^5,7^. Furthermore, coastal zones comprise multiple land-to-sea environmental gradients, including a range of salinities: freshwater-brackish-marine, leading to a large diversity of habitats^5,8^.

The Baltic Sea and the adjacent Kattegat-Skagerrak (hence “Baltic Sea area” Fig. 1a) are among the best-studied coastal sea areas^9^. Recent formation (8000 years ago^10^), pronounced environmental gradients, and narrow connecting straits make the Baltic Sea a useful model for studying eco-evolutionary processes driven by dispersal and adaptation^11^. In addition, its early exposure to anthropogenic pressures, especially eutrophication, can foreshadow the processes awaiting other coastal regions^12^. The Baltic Sea area also has a long record of microbial research, with the earliest data coming from the 19th century^13^. Regular microscopy-based phytoplankton monitoring has been run by multiple countries in the region for decades^14^ and since the 1980s in Sweden^15^. More recently, the Baltic Sea area was one of the first brackish waters surveyed using DNA metabarcoding^15–17^ and metagenomics^18,19^. Nevertheless, most of the unknown diversity in the Baltic Sea area is believed to be microbial^20^. Up until now, research has mainly focused on specific groups of phytoplankton and metazooplankton^20^, with much of the microbial diversity missing from the picture.

**Fig. 1.**
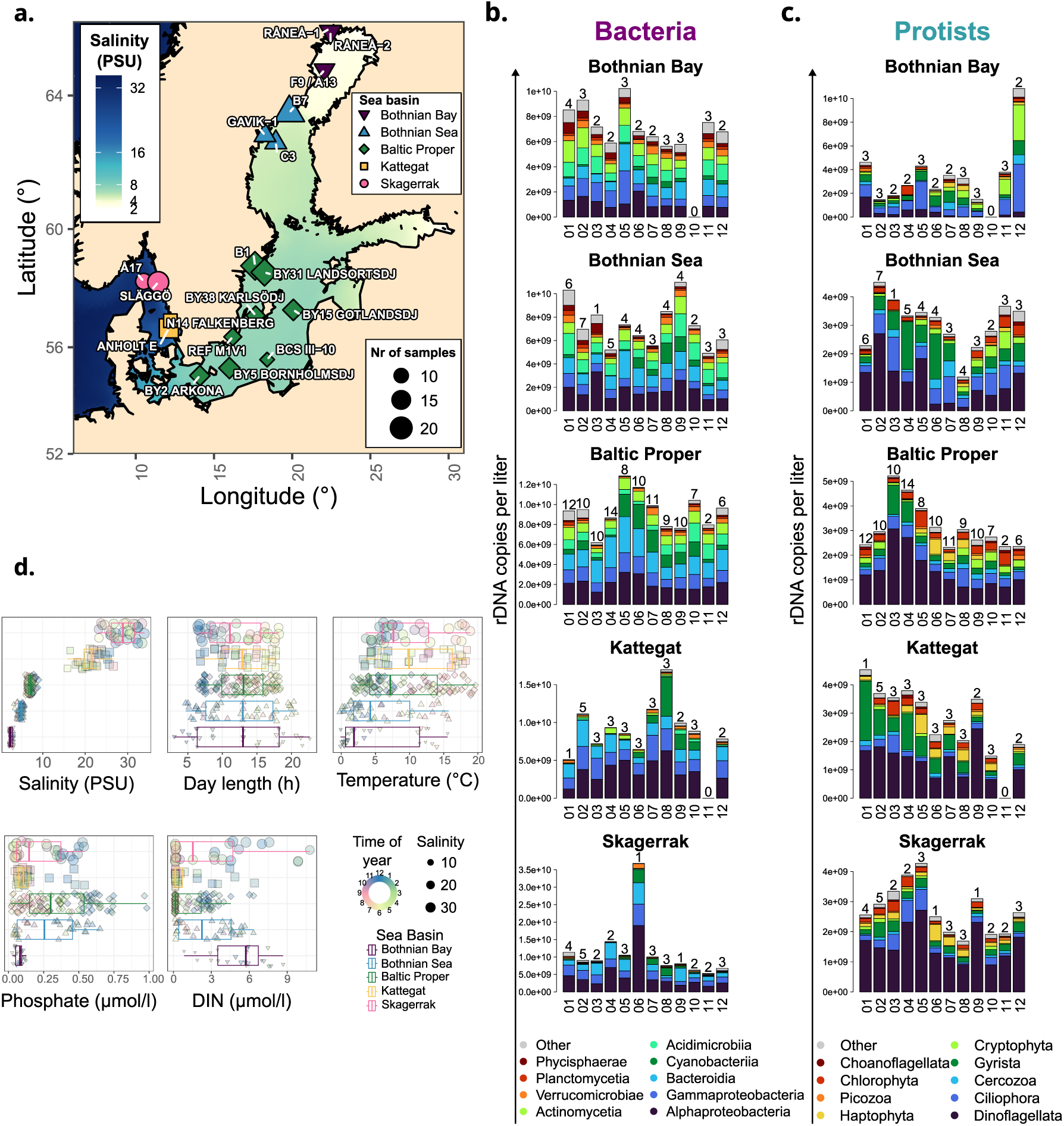
Spatiotemporal changes in abundance of major microbial taxa, 2019-2020. **a.** A map of 18 sampling locations labeled by the name of the station and size-coded by the number of samples collected over a period of 13 months (January 2019 to February2020). **b-c.** The numbers of rRNA marker gene copies per liter annotated to **b.** bacterial classes and **c.** protist subdivisions averaged over each month and Baltic Sea area basin (as marked in **a.**). The values are based on the number of reads relative to spike-in reads and the amount of spike-in added per volume of sample. Only the classes/subdivisions which, on average, corresponded to >0.01 of reads across samples are shown. Above each bar, the number of samples collected in the respective month and basin is given. Note: Two samples (January in Bothnian Bay and September in Bothnian Sea) were removed for protists due to spike-in detection failure. **d.** The distribution of measurements of major physicochemical and geographic parameters for each basin. DIN - dissolved inorganic nitrogen.

Globally, salinity is among the strongest factors structuring bacterial^21–24^ and protist^25^ communities. However, the importance of salinity has largely been inferred from the divide between freshwater and marine environments^26–28^. Previous research shows that brackish waters host unique bacterial species^19,29^. Both microscopy- and DNA-based surveys in the Baltic Sea area suggest pronounced shifts in surface water microbial community composition (beta diversity) along the horizontal salinity gradient^16,17,30–32^. Moreover, bacterial communities in the Baltic Sea change more with salinity than between winter and summer^33^. More generally, results obtained across steep salinity gradients in Antarctica suggested stronger environmental filtering among planktonic bacteria than protists^34,35^, while studies in other environmental contexts came to the opposite conclusion^36–39^. Whether bacterial communities are more affected by salinity in particular, or generally more/less environmentally filtered than protist ones, is yet to be determined.

The number of aquatic animal species tends to reach a minimum within the brackish spectrum across salinity gradients^20,40^. For phytoplankton, patterns of both a maximum^30^ and a minimum^31^ of species richness (alpha diversity) within the brackish salinity spectrum, as well as an increase along the salinity gradient^32^, have been reported based on microscopic assessments. DNA metabarcoding transect surveys of plankton showed no clear patterns for bacteria^16,17,33^ nor major eukaryotic taxa^17^. At the same time, bacterial alpha diversity has been observed to be lower in winter than in summer in the Baltic Sea^33,41^, consistently with patterns observed in marine environments^42–44^. In contrast, microscopy-based analysis of phytoplankton alpha diversity has shown different seasonal patterns in low and high salinities in the Baltic Sea area^32^. Little is known about the factors driving the seasonal patterns or confounding diversity and community composition changes along the salinity gradient.

Here, we present the first in-depth comparative analysis of bacterial and protist diversity across spatiotemporal scales in the Baltic Sea area and across the corresponding environmental gradients. We use a recently published DNA metabarcoding dataset^45^ and new data released here, which together span two complete calendar years and include 398 samples. We show how protists and bacteria differ in their ecological sensitivity to salinity and analyze drivers of taxonomic richness. Finally, we propose an explanation of the contrasting bacterial and protist diversity patterns based on fundamental physiological and ecological differences between the groups.

## Results and discussion

### Processing of metabarcoding data for ecologically relevant metabarcoding-based analyses

We based the first part of our analyses on a publicly available 16S and 18S metabarcoding dataset, covering broad spatiotemporal scales and environmental gradients regularly sampled throughout a year^45^. After filtering procedures (see Methods for details), we used the data from 246 independent samples. They came from 18 locations, extending from the Bothnian Bay (minimum salinity = 2.0 PSU) to the Skagerrak (maximum salinity = 33.92 PSU), each location sampled at least six times between January 2019 and February 2020 (Fig. 1a).

From the 16S amplicon sequence variants (ASVs), we chose only those classified as *Bacteria* and excluded plastids and mitochondria. From the 18S metabarcoding results, we excluded ASVs classified as animals (*Metazoa*), fungi, land plants (*Embryophyceae*), and selected macroalgae. We thus treated the remaining 16S ASVs as bacteria and 18S ASVs as protists.

We clustered the ASVs by simultaneous genetic and distribution similarity, using distribution-based operational taxonomic unit-calling (dbOTU^46^; we thus call those taxonomic units dbOTUs). This method groups variants corresponding to different marker gene copies within one organism and other ecologically indistinguishable intra-species variation. Therefore, dbOTUs should correspond to ecologically coherent lineages unless the sequenced regions are identical between organisms with distinct ecologies. More than half of dbOTUs were annotated to at least family level for both bacteria and protists (Supplementary Fig. S1c-d). More than half of the reads were annotated to at least genus or family level for bacteria and protists, respectively (Supplementary Fig. S1e-f).

Finally, a controlled amount of synthetic DNA amplifiable by the metabarcoding primers, i.e., spike-in, was added to the samples. This approach aims to obtain absolute values (copy number per volume sample) for the marker gene (16S/18S ribosomal DNA (rDNA)), which better estimates absolute abundance than relative counts^47^. Thus, it can uncover blooming and succession patterns obscured from relative abundance analysis^48^. At the same time, the spike-in approach is highly sensitive to differences in sample preparation and amplification^47^. This can lead to noisier results than relative abundance, especially in environmental monitoring, as samples are often collected by multiple researchers working in variable conditions.

### Regionally distinct seasonal dynamics of major bacterial and protist taxa

Major bacterial (Fig. 1b) and protist (Fig. 1c) clades displayed distinct seasonal abundance dynamics. Both their abundance and seasonal patterns differed between the Kattegat-Skagerrak, and the three basins of the Baltic Sea included in this study (Baltic Proper, Bothnian Sea, and Bothnian Bay).

We observed an increased abundance of cyanobacteria as early as May in the Baltic Proper (Fig. 1b), consistent with microscopic observations^49^. The increase in cyanobacterial abundance was smaller and peaked later moving northwards within the Baltic Sea. In the Baltic Proper, both dinoflagellates and *Gyrista* (a protist subdivision that includes diatoms), peaked in abundance in March (Fig. 1c), as observed using microscopic methods^49,50^. The dynamics of *Dinoflagellata* and *Gyrista* differed vastly between basins. However, in all cases except for Bothnian Bay, those groups increased in abundance in the first half of the year. Furthermore, in the Kattegat and the Skagerrak, there was an additional dinoflagellate peak in September, corresponding to autumn blooms. *Haptophyta* generally displayed increased abundance between May and August. *Cryptophyta* were abundant in the Baltic Sea, with higher abundance in the second half of the year than in the first.

We also observed geographic differences in abundance and seasonal dynamics of major heterotrophic bacteria. *Bacteroidia* (Fig. 1b) peaked in abundance in May across the Baltic Sea, consistent with previous reports^41,51^, but displayed less clear seasonal dynamics in the Skagerrak and the Kattegat. *Actinomycetia* and *Acidimicrobiia* were abundant in the Baltic Sea but not in the Kattegat-Skagerrak, consistent with *Actinobacteria* abundance patterns from transect-based studies^16–18^. Additionally, *Acidimicrobiia* decreased in abundance between May and July, a dynamic that was more pronounced moving southwards in the Baltic Sea. Finally, contrary to previous results from a near-coast station^41,51^, *Alphaproteobacteria*, and *Gammaproteobacteria* comprised a stable proportion of the whole community throughout the year in all basins but the Kattegat (Supplementary Fig. S2a). Nevertheless, consistent with metabarcoding^16^ and metagenomic transects^18^, *Alphaproteobacteria* and *Gammaproteobacteria* were more abundant in the Kattegat-Skagerrak than in the Baltic Sea.

Overall, we observed geographic differences in both total abundance and seasonal dynamics of major bacterial and protist taxa. There were some latitudinal shifts, e.g., for cyanobacteria and *Acidimicrobiia*, and apparent differences between Kattegat-Skagerrak and the Baltic Sea. Compared to relative abundance-based analysis, spike-in correction added ecologically relevant information, as best exemplified by the March dinoflagellate bloom in the Baltic Proper (Fig. 1c) being indistinguishable based on the relative abundance of *Dinoflagellata* (Supplementary Fig. S2b). Still, the spike-in approach entailed substantial random noise, especially in less sampled basins (Fig. 1b-c) and relative abundances better captured gradual succession patterns, especially for bacteria (Supplementary Fig. S2a). Thus, the best normalization approach depends on the goals of the analysis and suitable statistical methods, at the very least until the consistency of absolute abundance quantification improves.

### Salinity has a dominant impact on bacterial but not protist community composition

The dissimilarities of bacterial and protist communities between the same pairs of samples were correlated (R2 = 0.58). However, the protist communities were generally less similar to each other than bacterial ones (Supplementary Fig. S3a), as has been reported for picoeukaryotes and bacteria in ocean surface waters52.

Principal Coordinate Analysis (PCoA) showed distinguishable geographic and seasonal structuring of bacterial and protist communities (Supplementary Fig. S4). To disentangle the environmental factors driving those structuring effects, we performed distance-based Redundancy Analysis (dbRDA). We focused on five fundamental environmental factors: salinity, temperature, day length (i.e., daytime length), dissolved inorganic nitrogen (DIN), and phosphate. These factors did not strongly correlate with each other (R2 < 0.4, Supplementary Fig. S5). The dbRDA-based variation partitioning showed that salinity could explain almost half of the variation in bacterial community composition (Fig. 2a). The effect of salinity on protist community composition was much smaller (13%), while temperature and nutrients were of greater relative importance than for bacteria (Fig. 2b). Importantly, these results contradict the proposed rule that the distribution of smaller organisms is less limited by environmental factors than that of larger organisms^53^, and in particular that bacteria are less environmentally filtered than protists^38^. At the same time, our results are consistent with a previous report of a stronger selection of bacterial than protist communities across a salinity gradient^35^. However, they do not support the generalization of the stronger selection to the effects of other environmental factors.

**Fig. 2.**
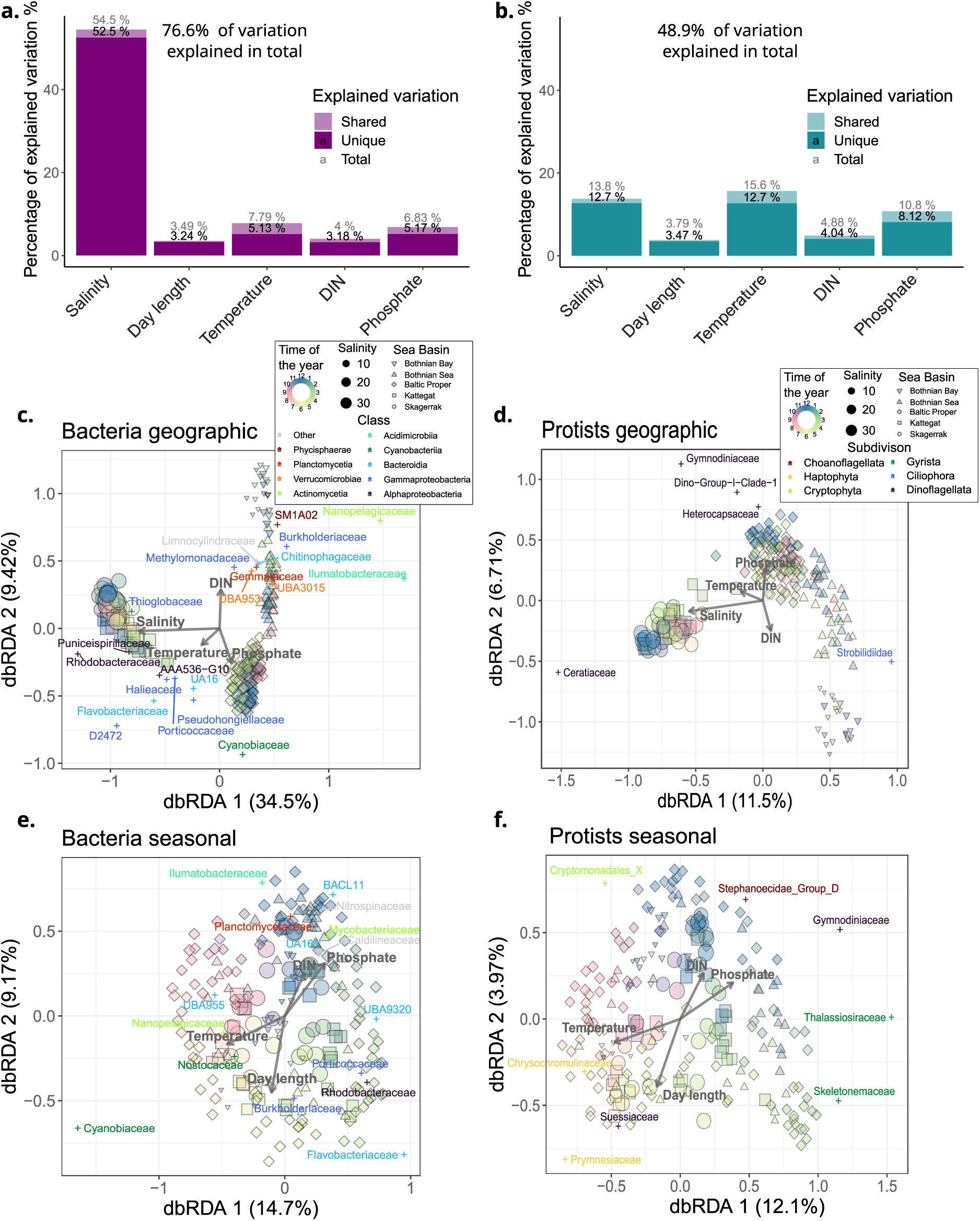
Environmental factors explaining geographic and seasonal community structuring and the most correlated families of bacteria and protists. **a-b.** Variance partitioning of bacterial (**a.**) and protist (**b.**) beta diversity, based on Bray-Curtis distances and distance-based Redundancy Analysis (dbRDA). **c-d**. Partial dbRDA conditioned on seasonality (sine and cosine of the day of the year), highlighting bacterial (**c.**) and protist (**d.**) geographic community structuring. **e-f**. Partial dbRDA conditioned on salinity and latitude, highlighting bacterial (**e.**) and protist (**f.**) seasonal community structuring. **c-f**. Weighted averages of families deviating (Bonferroni corrected *P* < 0.05) were placed on the plot, with values rescaled by a factor of 0.5. Percentages of variation explained by each dbRDA component are given in brackets by the axes labels. DIN - dissolved inorganic nitrogen. Salinity is given in practical salinity units (PSU).

Nevertheless, for both bacterial and protist communities, there was a pronounced salinity divide between the Baltic Sea and the Kattegat-Skagerrak and a more gradual latitudinal shift within the Baltic Sea (Fig 2c-d). The latter corresponded to decreasing salinity and phosphate-to-nitrogen ratios with increasing latitude (Fig. 1d). We connected those spatial changes in beta-diversity, calculated from ASV-based dissimilarities, with high relative abundance of specific microbial families (Fig 2c-d). For example, typically freshwater *Nanopelagicaceae* (acI *Actinobacteria*)^54^ and *Limnocylindraceae* (*Chloroflexi*)^55^ were associated with the lowest salinity samples, while commonly marine *Rhodobacteraceae^56^* and *Thioglobaceae^57^* were at the other end of the salinity spectrum (Fig 2c).

Moreover, both bacterial and protist communities underwent gradual community composition changes throughout the year in all the analyzed basins of the Baltic Sea area (Fig. 2e-f, Supplementary Fig. S6). Diverse groups significantly contributed to the seasonal patterns in beta diversity, ranging from known seasonally blooming diatoms *Thalassiosiraceae* and *Skeletonemaceae^58^* to multiple uncharacterized *Bacteroides* families. In all the basins, picocyanobacteria (*Cyanobiaceae*) were associated with high temperatures, while BACL11 (*Bacterioides^19^*) were typical of winter conditions (Fig. 2e, Supplementary Fig. S6a).

### Interannual stability of microbial community composition

To assess the interannual stability of the community composition patterns we observed, we performed metabarcoding on stored DNA extracted from samples collected between 2015 and 2017 (Fig, 5a-b). The new dataset, corresponding to 143 samples, covered the southern range of the 2019-2020 dataset (Fig. 1a, Fig. 3a). Similarly to the 2019-2020 dataset, it mainly corresponds to a 13-month long time-series, between February 2016 and March 2017, with a few extra samples from 2015 from one station (Släggö, Fig. 3a). There were minor differences in sample processing between the two datasets, most notable being: DNA extraction kit, different Illumina sequencing systems, and lack of spike-in sequences in the 2015-2017 dataset (see Methods for more details).

**Fig. 3.**
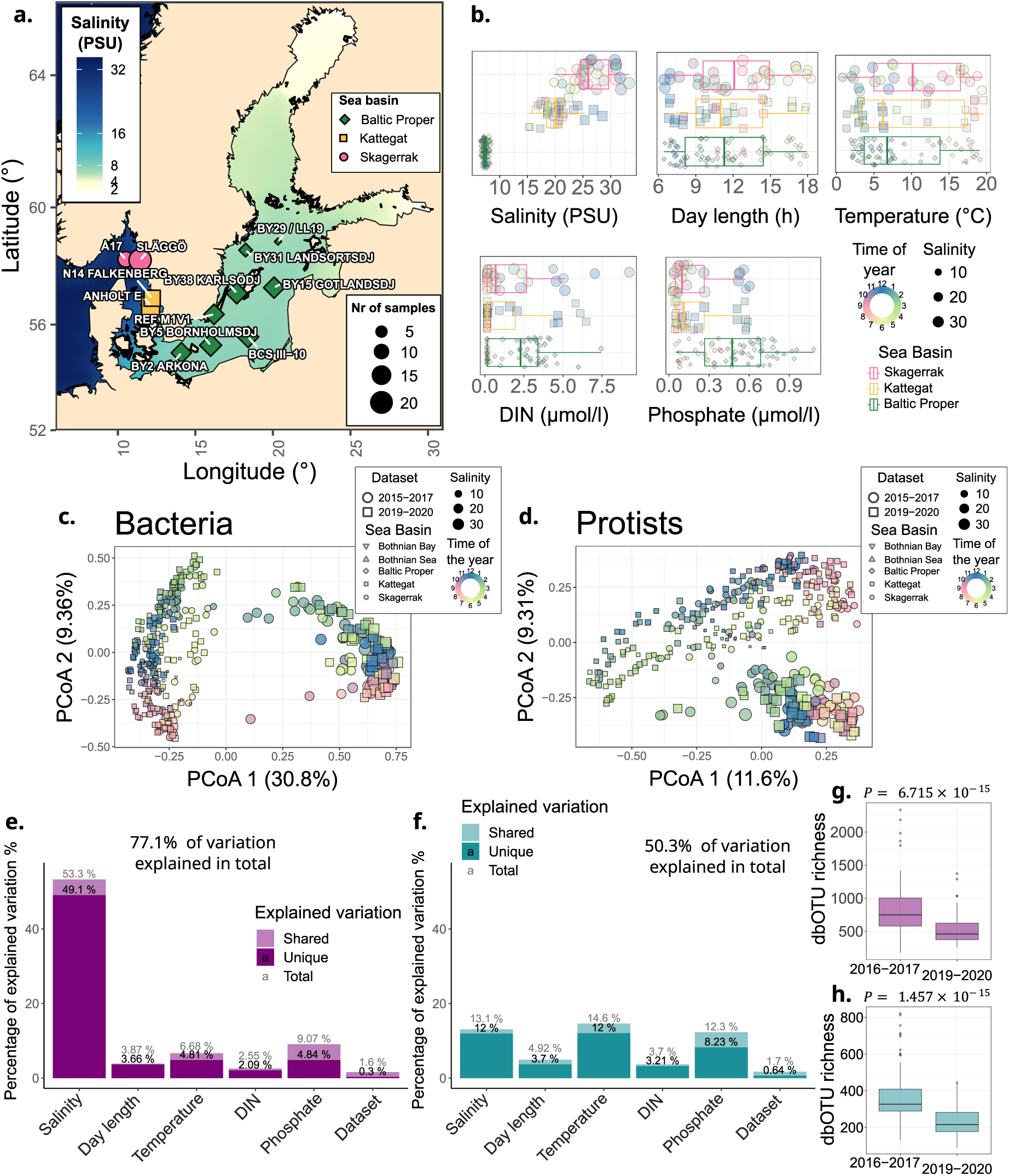
Bacterial and protist diversity across datasets collected in 2015-2017 and in 2019-2020. **a.** A map of 12 sampling locations labeled by the number of samples collected between August 2015 and March 2017 (the 2015-2017 dataset), with all the 2015 samples coming from station Släggö in Skagerrak **b.** The distribution of measurements of major physicochemical and geographic parameters for each basin in the 2015-2017 dataset. **c-d**. PCoA based on Bray-Curtis dissimilarities of bacterial (**c.**) and protist (**d.**) communities across the 2015-2017 and 2019-2020 datasets. **e-f.** Variance partitioning of bacterial (**e.**) and protist (**f.**) diversity, based on Bray-Curtis distances and dbRDA. Dataset stands for a dummy variable differentiating the samples collected in 2015-2017 and 2019-2020. **g-h.** dbOTU richness of bacteria (**g.**) and protists (**h.**) in the two datasets. Samples of both datasets were rarefied to the same number of reads. Only dbOTU richness values for stations present in both datasets have been used. Moreover, samples were chosen in pairs, from the same station and the same month of the year but different datasets. Sample pairs with the lowest distance in terms of the day in a calendar year were chosen. *P-*values obtained using the pairwise Wilcoxon (signed-rank) test are given above each plot. DIN - dissolved inorganic nitrogen. dbOTU - distribution-based operational taxonomic unit.

Overall, the community composition in the two datasets differed minimally, following the spatial and seasonal environmental gradients analogously (Fig. 3c-f). This apparent interannual stability of community composition is consistent with previous observations from the Baltic Sea^51,59^ and other waters^39^. Finally, the similarity of communities between the two datasets shows that the technical differences had little impact on the observed general community composition (Fig. 3e-f).

Only a two-year break separated the datasets. The seasonal patterns likely change over longer time scales, as, e.g., climate change has altered the phytoplankton bloom dynamics in the Baltic Sea over decadal scales^60^. As climate change accelerates^61^ and nutrient inputs are being decreased in the Baltic Sea^62^, the microbial communities will surely undergo long-term changes. The introduction of invasive species can also alter microbial community dynamics^63^. A reproducible monitoring scheme encompassing diverse taxa, achievable with DNA-based methods, is crucial for understanding and informed responses to unforeseen environmental changes.

In contrast to community composition, alpha diversity differed significantly (*P* < 10^-14^ in both cases) for both bacteria and protists (Fig. 3g-h). We doubt the increased diversity in 2015-2017 dataset, amounting to hundreds of dbOTUs on average, represented an actual biological change. Instead, it likely originated from the technical differences between the datasets. We used rarefaction for alpha-diversity estimates, as it performs best in removing sequencing depth biases^64^ (expected due to different Illumina sequencing systems used for the datasets). Nevertheless, in both datasets, rarefaction curves were saturated, suggesting that extra reads would not change alpha diversity much (Supplementary Fig. S7). Thus, another technical difference, remaining to be identified, likely caused the dbOTU difference between datasets. Ultimately, our results suggest more caution is needed when comparing alpha diversity than beta diversity across datasets. As a consequence, in subsequent alpha diversity analyses, we focused solely on the 2019-2020 dataset because of its larger geographic scope.

### Seasonal vertical mixing brings deep-water bacteria to the surface and increases bacterial alpha diversity

We assessed the changes in alpha diversity along the salinity gradient, for both bacteria and protists, following the methodology used to argue for or against maximum^17,30^ and minimum^31^ protist diversity along the gradient (Fig. 4a-b). The horohalinicum, i.e. midrange salinities (5-8 PSU), displayed lower bacterial richness than in less (but not more) saline waters (Fig. 4a). The minimum in bacterial richness was at 15 PSU. However, the minimum estimation was likely confounded by seasonal changes in salinity within Kattegat, as there was clear seasonal variation in bacterial richness overshadowing the geographic differences (Fig. 4a, c).

**Fig. 4.**
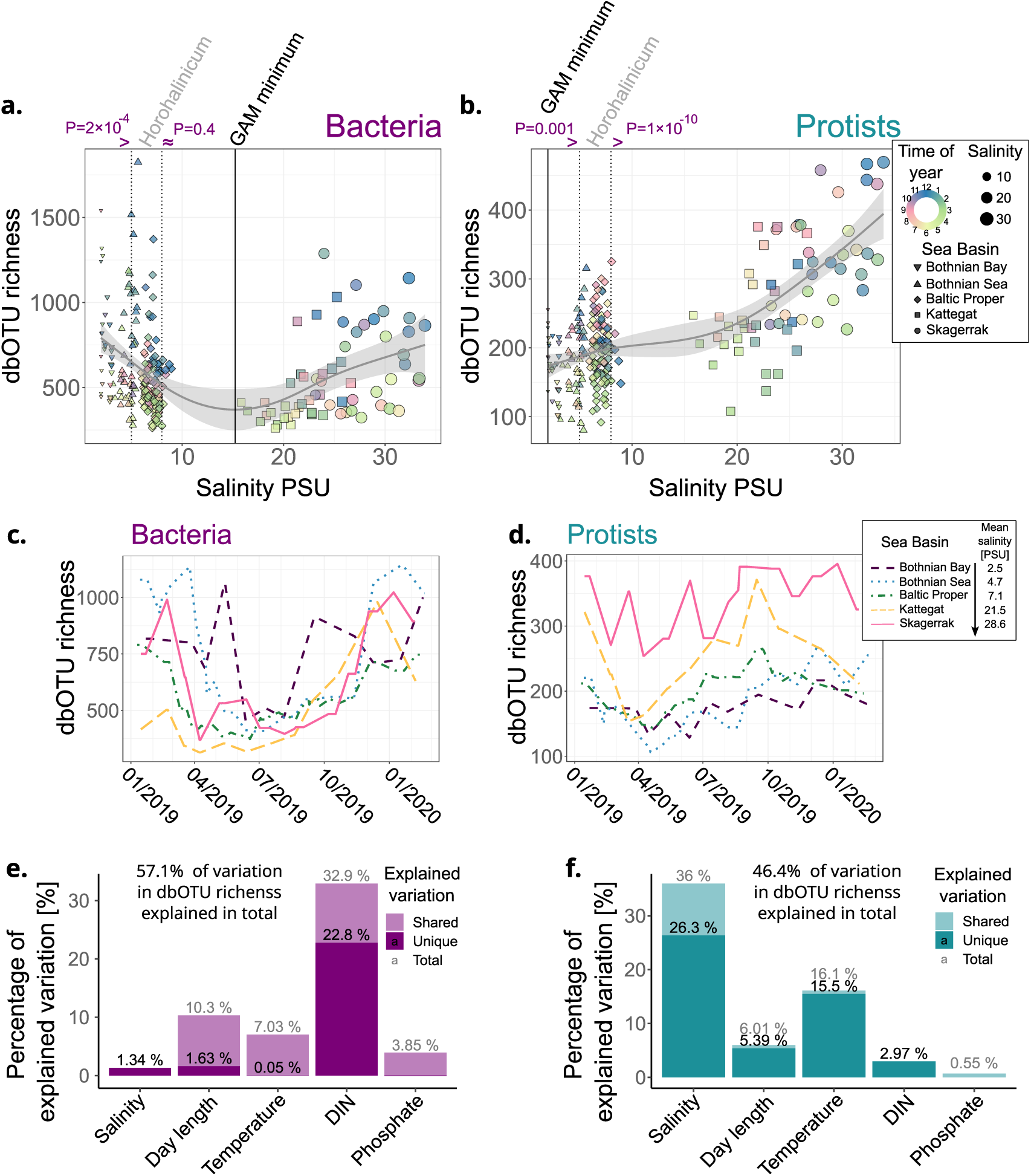
Bacterial and protist richness. Mean number of bacterial (**a.**) and protist (**b.**) dbOTUs found across 100 refraction iterations (dbOTU richness). The fit corresponds to smoothened generalized additive models (GAMs). The dotted vertical lines correspond to boundaries of horohalinicum (5-8 PSU). The colored (in)equality signs over the dotted lines mark the significant differences of dbOTU richness between horohalinicum and lower/higher salinities, and the P-values from Wilcoxon rank-sum tests are written above. The solid vertical lines mark salinities corresponding to minimum richness in the GAMs. The same legend applies to panels a-b. **c-d.** Rolling means of bacterial (**c.**) and protist (**d.**) richness for the distinguished Baltic Sea area basins. The richness values were averaged for measurements two weeks before and after each date. The same legend applies to panels c-d. **e-f**. Variance partitioning of bacterial (**e.**) and protist (**d.**) dbOTU richness.

Across geographic regions, bacterial alpha diversity rapidly declined in spring and increased again in autumn (Fig. 4c). Out of the assessed environmental factors, DIN explained the largest part of its variation (Fig. 4e). Moreover, DIN followed similar seasonal patterns of rapid decline and increase (Supplementary Fig. S8a) and correlated with seasonal rather than geographic variation in dbOTU richness (Supplementary Fig. S8b).

However, the spring depletion of nitrogen in the Baltic Sea is closely connected to the emergence of the summer thermocline and thus reduced vertical mixing^65^. Winter vertical mixing has been linked to bacterial alpha diversity and similarity in cytometric properties across the water column in a marine time series^66^. Additionally, some members of typically deepwater clades are rare taxa in surface waters with abundance maxima in winter^67^. The influx of deepwater bacteria has also been suggested to explain the higher diversity of bacteria in winter than in summer in the Baltic Sea, with winter communities being less structured by depth^33^. However, these patterns may originate from similar conditions across the water column, leading to related bacteria thriving. Evidence that deepwater bacteria actually influx to the surface in winter is missing.

Therefore, we connected our results with a previously published metabarcoding dataset spanning different depths across a winter and a summer transect in the Baltic Sea^33^, from here on referred to as the “transect-based dataset”. We matched the sequences representative of the OTUs (operational taxonomic units) distinguished in the transect-based dataset to ASVs representative of our dbOTUs (see Methods for details). Despite substantial methodological differences, we found a match for 36.6% of the OTUs and 16.1% of dbOTUs at a 99% nucleotide identity threshold^68^.

We further chose from the transect-based dataset only the summer (July) samples, corresponding to the time of strong thermocline. Among those, we categorized OTUs as found only in deep (>10m depth) or in surface (≤10m depth) waters, or in both. We assessed how many OTUs from each category matched the dbOTUs from winter (December-February) and summer (June-August) months (Fig. 5a). We excluded Bothnian Bay from the analysis since there were no deepwater samples from this region in the transect-based dataset. Among the deepwater OTUs, significantly more (*P* < 10^-75^) were found in winter than in summer despite originating from summer samples. In contrast, more OTUs found in both deep and surface water were detected in summer (*P* < 10^-8^), while among the surface OTUs, there was no significant difference between the seasons (*P* = 0.67). Ultimately, these results suggest a substantial proportion of taxa found only in deep waters in summer can be found at the surface during winter.

**Fig. 5.**
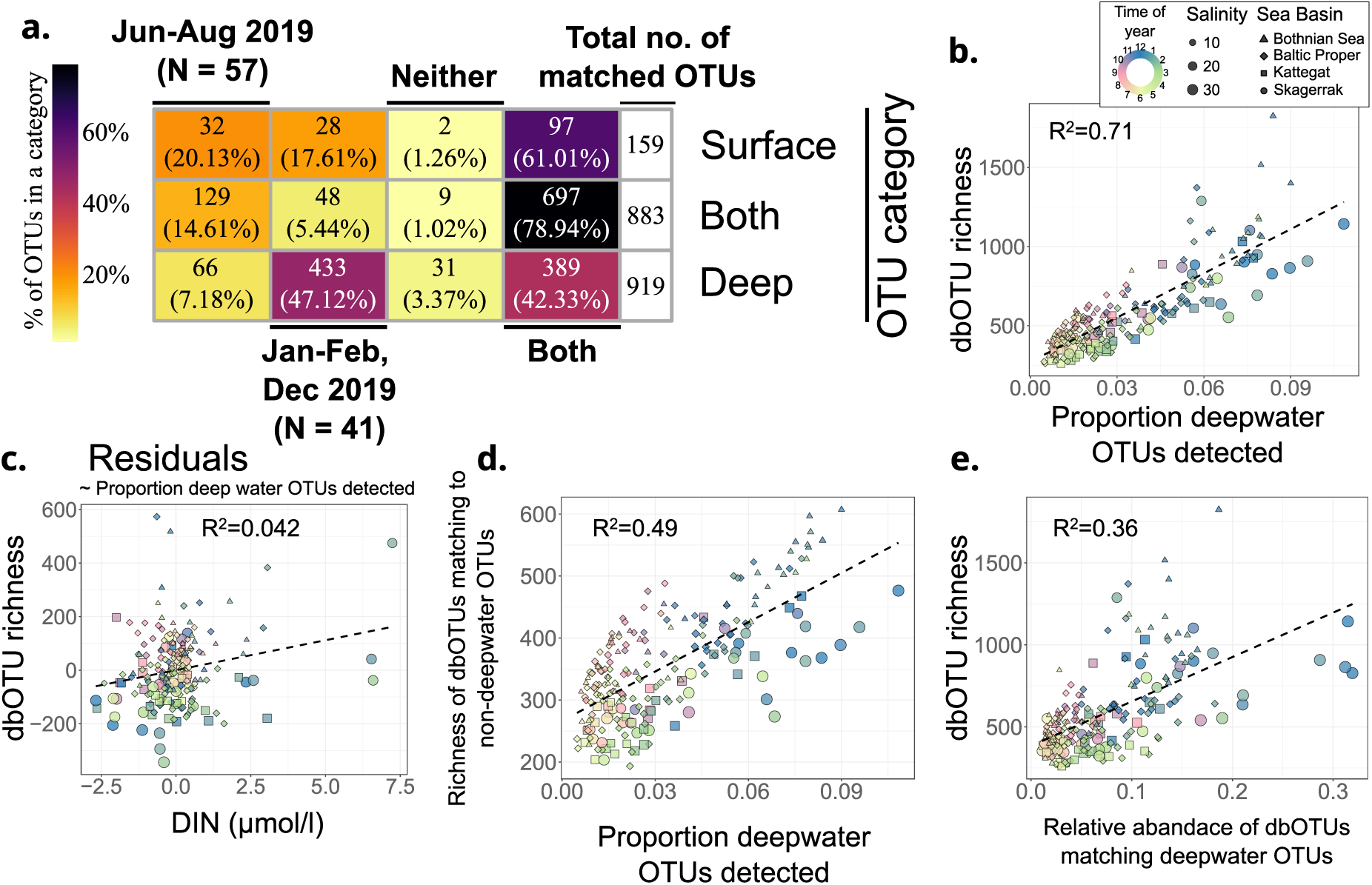
Bacterial alpha diversity and the proportion of deepwater OTUs detected. **a**. Number of OTUs from a previous, transect-based study^33^ matched to dbOTUs found in winter and summer samples or in neither or both seasons. The OTUs are categorized by being found exclusively in deep water (> 10m depth) samples, exclusively in surface waters, or in both, in the original study. Only OTUs matching to any dbOTUs from this study are counted. **b.** Proportion of deepwater OTUs (including ones without a match in the 2019-2020 dataset) having a match to a dbOTU from a sample vs. bacterial dbOTU richness. **c.** The residuals of DIN vs. dbOTU richness from models correlating the factors to the proportion of deepwater OTUs detected, corresponding to partitioning out the unvisualized parameter. **d.** Proportion of all deepwater OTUs vs. richness of bacterial dbOTUs matching to non-deepwater-specific OTUs (“surface” and “both” categories). **e.** Relative abundance of all dbOTUs matching a deep water OTU vs bacterial dbOTU richness. Bothnian Bay was excluded from the plots since there were no deepwater samples taken in this region in the transect-based study^33^. The same legend applies to figures b-e. DIN - dissolved inorganic nitrogen.

The proportion of detected deepwater OTUs in a sample correlated strongly with dbOTU richness, with a clear seasonal pattern (Fig. 5b). In contrast, the proportion of detected non-deepwater-specfic OTUs (‘surface’ and ‘both’ from Fig. 5a) showed a weaker correlation with richness (Supplementary Fig. S8c), and did not follow the proportion of deepwater OTUs matched (Supplementary Fig. S8d). Thus, it was not the detection biases that underpinned the correlation between the proportion of deepwater OTUs and richness. Moreover, after partitioning out the correlations with deepwater OTUs, DIN no longer correlated with dbOTU richness (Fig. 5c). This relation was not bilateral, as the proportion of deepwater OTUs still correlated with dbOTU richness after partitioning out DIN (Supplementary Fig. S8e). Therefore, the correlation between DIN and bacterial diversity comes from a common confounder: seasonal convective water mixing.

However, the richness of dbOTUs matching to non-deepwater-specific OTUs still correlated with the proportion of deepwater OTUs (Fig. 5d). Simultaneously, we detected more non-deepwater-specific OTUs in summer than in winter (Fig. 5a). Therefore, the same taxa were more persistently detected across the samples in winter. This could not be explained by a few dominant taxa pushing others below detection limits in summer, as the correlation remained after partitioning out Pielou’s evenness index^69^ (Supplementary Fig. S8f). Thus, some consequences of vertical mixing or parallel processes, e.g., lower irradiation^44,70,71^, lead to a more persistent or geographically widespread presence of the same taxa in the surface waters in winter than in summer.

In winter, deepwater taxa not only increased alpha diversity but also comprised a substantial part of the surface water bacterial communities, up to >30% in near-marine conditions (in Skagerrak; Fig. 5e). In contrast, surface water-specific taxa never reached cumulative relative abundance higher than 0.1 (Supplementary Fig. S8g), while taxa found in both surface and deep waters always corresponded to more than half of the reads (Supplementary Fig. S8h). Thus, the ‘both’ category likely includes abundant surface water taxa found also in deeper waters in summer due to sinking.

With cumulative relative abundance commonly above 0.1 in winter (Fig. 5e), the deepwater taxa likely impact the surface water ecosystem, if not through metabolic activity, at least by providing an alternative food source. Even if the deepwater taxa can survive only briefly at the surface, they can still substantially impact longer-term community dynamics^72^. A further investigation of the consequences of these community coalescence events is needed.

Ultimately, the higher mean bacterial richness in the least saline Bothnian Sea and Bothnian Bay may well be explained by longer vertical mixing periods due to colder climates. In summer, when mixing ceased, richness dropped to similar levels across basins (Fig. 4c), consistent with previous findings reporting no salinity-dependent pattern of bacterial alpha diversity^16,17,33^.

### Higher protist alpha diversity in near-marine salinities

In contrast to the situation for bacteria, protist richness displayed a clear salinity-dependent pattern, increasing with salinity (Fig. 4b; linear fit yields R^2^=0.44). This is consistent with a recent microscopy-based study^32^. However, unlike in the microscopy-based study, richness in the near-marine Kattegat-Skagerrak was higher than in the Baltic Sea across the year (Fig. 4d).

Similarly to our observations on protists, metazooplankton and fish alpha diversity in the Kattegat-Skagerrak is higher than in the Baltic Sea due to many marine species in thriving in higher but not lower brackish salinities, outweighing the presence of freshwater species in lower salinities^20^. The Baltic fauna comes mainly from a limited number of local adaptations to brackish conditions following the last glaciation period^11,20^. In contrast, for Baltic Sea bacteria, we recently showed evidence against widespread occurrence of such local adaptation, but rather colonization by a global brackish microbiota^29^. Consistently, this (Fig. 4a, c) and previous studies^16,17,33^ showed no evidence of decreased bacterial diversity in the Baltic Sea compared to Kattegat-Skagerrak. Whether the observed protist diversity pattern originates from a selected group of locally adapted species, as is the case for animals, is yet to be determined.

Temperature was the second factor most strongly associated with protist alpha diversity (Fig. 4f), and in most basins, dbOTU richness increased from March to September (Fig. 4d). Temperature was also the factor best explaining protist beta diversity (Fig. 2b). Therefore, our results suggest that both common and rare protists might be more susceptible to the direct effects of climate change than bacteria.

### Protist dbOTUs more often occur at both high and low brackish salinities

The salinity gradient in the Baltic Sea area includes a rapid shift across the Danish straits. Consequently, the data we used contained geographically proximate samples from lower (<9 PSU) and higher (>15 PSU) brackish salinities (Fig. 1a, Fig. 3a). The above analyses showed this salinity barrier to have the strongest structuring effect on the microbial communities, though less so for protists than for bacteria (Fig. 2 and 3). On the other hand, protist but not bacterial alpha diversity was higher on the more saline side of the barrier (Fig. 4).

We further investigated whether bacterial or protist dbOTUs were generally less likely to cross this salinity barrier between lower and higher salinities. To minimize the effects of other environmental filtering effects (especially climate), we chose the same number of stations and samples on both sides of the salinity barrier (see Methods for details). Additionally, we based further analyses on the 2019-2020 dataset, which allowed us to use spike-in normalized counts as an abundance metric comparable between bacteria and protists. The proportion of protist dbOTUs crossing the salinity barrier across the subsequently selected samples was significantly higher than that of bacterial dbOTUs (Fig. 6a).

**Fig. 6.**
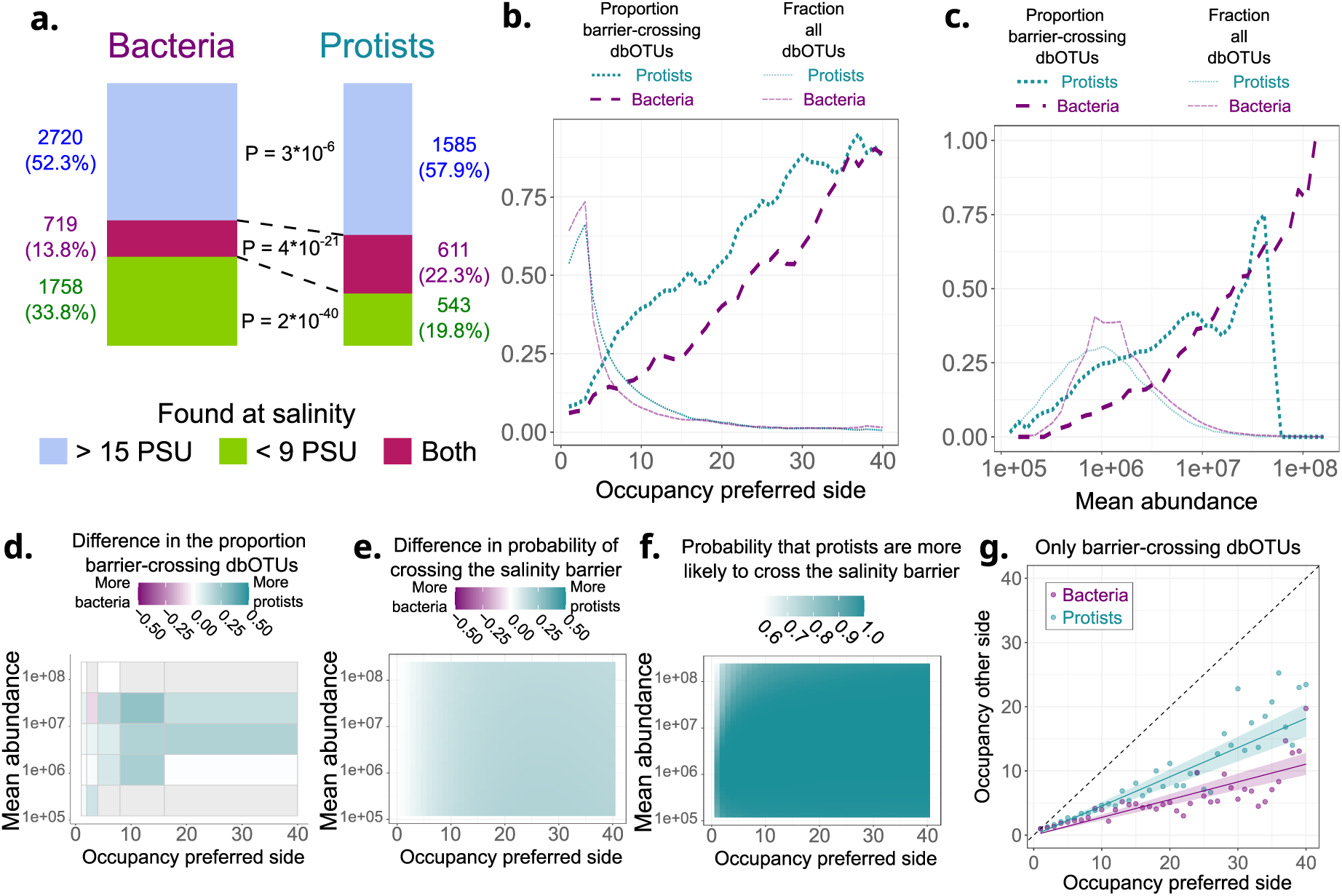
Proportion and probability of bacteria and protists (being) found in both high (>15 PSU) and low (<9 PSU) brackish salinity. **a.** The number and proportion of bacterial and protists dbOTUs found in high (>15 PSU), low (<9 PSU) brackish salinity, and across the salinity barrier. **b-c.** The proportion of barrier crossing dbOTUs for each value of maximal occupancy at one side of the salinity barrier (**b.**) and across forty bins of mean abundance (**c.**). The proportion of all dbOTUs that lie within each bin is also given. The values are rolling means across +/- two bins. **d.** The difference in the proportion of protist and bacterial barrier crossing dbOTUs. Grey tiles correspond to the values across which either no protist or no bacterial dbOTUs were found. **e-g.** Results from Bayesian modeling. **e.** The mean difference in the predicted probability of a protist and a bacterial dbOTU crossing the salinity barrier. **f.** The probability (proportion of MCMC iterations) that protists are more likely to cross the salinity barrier. **g.** The occupancy on the less occupied side of the salinity barrier as a function of the occupancy on the more occupied side. The points correspond to means of the experimental values. The lines correspond to the values predicted from the model with 95% confidence intervals. The dashed black line is the identity line. For all the analyses, only the four stations with salinity >15 PSU, and four out of the most saline stations with salinity <9 PSU are chosen. The data for each station is downsampled to the same number of observations. During downsampling, samples closest in time to the samples from the least sampled station were chosen. Only observations from 2019-2020 are included. The abundance values correspond to numbers of rDNA copies per liter, based on the spike-in normalization, and are always given on a logarithmic scale. “Occupancy preferred side” and “occupancy other side” refer to the occupancy at the side with highest and lowest occupancy, respectively.

Still, the observed proportion of barrier-crossing bacteria and protists may be affected by detection biases coming from differences in relative abundance, length of vegetative season, and environmental filtering by factors other than salinity. Bacteria were, on average, more abundant in terms of rDNA gene marker copy numbers (Supplementary Fig. S9a). As protists tend to have higher rDNA copy numbers per cell than bacteria^73^, this difference likely translates to an even larger difference in cell numbers, a more relevant abundance measure in the context of dispersal capabilities and detection probability. Meanwhile, on average, protists were present in more samples on their preferred side (side with higher occupancy) of the salinity barrier (Supplementary Fig. S9b). Overall, similarly widespread bacteria were more abundant than protists (Supplementary Fig. S9c-f). Still, across the occupancy values, a higher proportion of protists crossed the salinity barrier (Fig. 6b). The same was true across the abundance values, apart from the most abundant outliers, which were barrier-crossing dbOTUs among bacteria but not protists (Fig. 6c). Generally, across different occupancy and abundance values, a higher proportion of protists crossed the salinity barrier (Fig. 6d, Supplementary Fig. S9g-h).

To confirm that protists are more likely to cross the salinity barrier, we build a Bayesian model using a Monte Carlo Markov Chain (MCMC) with Just Another Gibbs Sampler^74^ (JAGS). The model, based on the Bernoulli distribution, calculates the probability of crossing a barrier from logistic regression from the maximal occupancy on one side of the barrier, geometric mean abundance, and whether a dbOTU is a protist or a bacterium. After model selection using the Watanabe-Akaike information criterion^75^, we chose a version of the model assuming the probability dependence on the other variables follows different functions for bacteria and protists (see Methods for details). The model predicted protists to be more likely to cross the salinity barrier across the occupancy and abundance values, both on average (Fig. 6e, Supplementary Fig. S9i-j) and in a majority of MCMC iterations (Fig. 6f). The probability that protists are more likely to cross the salinity barrier (Fig. 6f) was consistently close to one except for dbOTUs with low occupancy and high abundance, in which case detection biases might outweigh the biological signal.

Finally, using a similar approach, we built a Bayesian model of total occupancy as a function of the occupancy on the more occupied side of the salinity barrier, based on the Poisson distribution. From this model, we calculated the predicted occupancy on the less occupied side of the salinity barrier and compared it to the experimental values (Fig. 6g). The model and the data showed that protists not only more often crossed the salinity barrier but also were, in general, more widespread in the less preferable salinity regime.

### Proposed explanation of the stronger ecological sensitivity to salinity among bacteria than protists

We propose that compartmentalization makes eukaryotic cells (thus, protists) more acclimatable than prokaryotic ones (thus, bacteria) to a different salinity regime (Fig. 7). In a compartmentalized cell, crucial metabolic processes take place across internal membranes. External ion concentrations and pH do not directly affect these processes. Notably, in eukaryotic cells, the proton motive force (PMF) used for ATP synthesis is maintained along the inner membranes of mitochondria and plastids. In contrast, prokaryotes usually produce PMF across the cellular membrane directly exposed to the external environment. While eukaryotic cytosolic pH is kept at stable, neutral levels^76^, marine and brackish waters are buffered and usually slightly alkaline (pH∼8)^27,77^. Thus, in the presence of sea salt, some transferred protons are consumed in reactions with (bi-)carbonates or hydroxide. Consequently, many marine bacteria preferably use sodium motive force (SMF), i.e., Na^+^-gradient across the membrane, for transmembrane transport, motility, and/or ATP production^18,27,78,79^. The competitive advantage of this strategy diminishes with decreasing salinity (in a logarithmic manner, assuming stable intracellular ion concentrations^80^), and so does likely the relative abundance of bacteria using it. Additionally, many protists have organelles with osmoregulating functions, especially the contractile vacuole^81^.

**Fig. 7.**
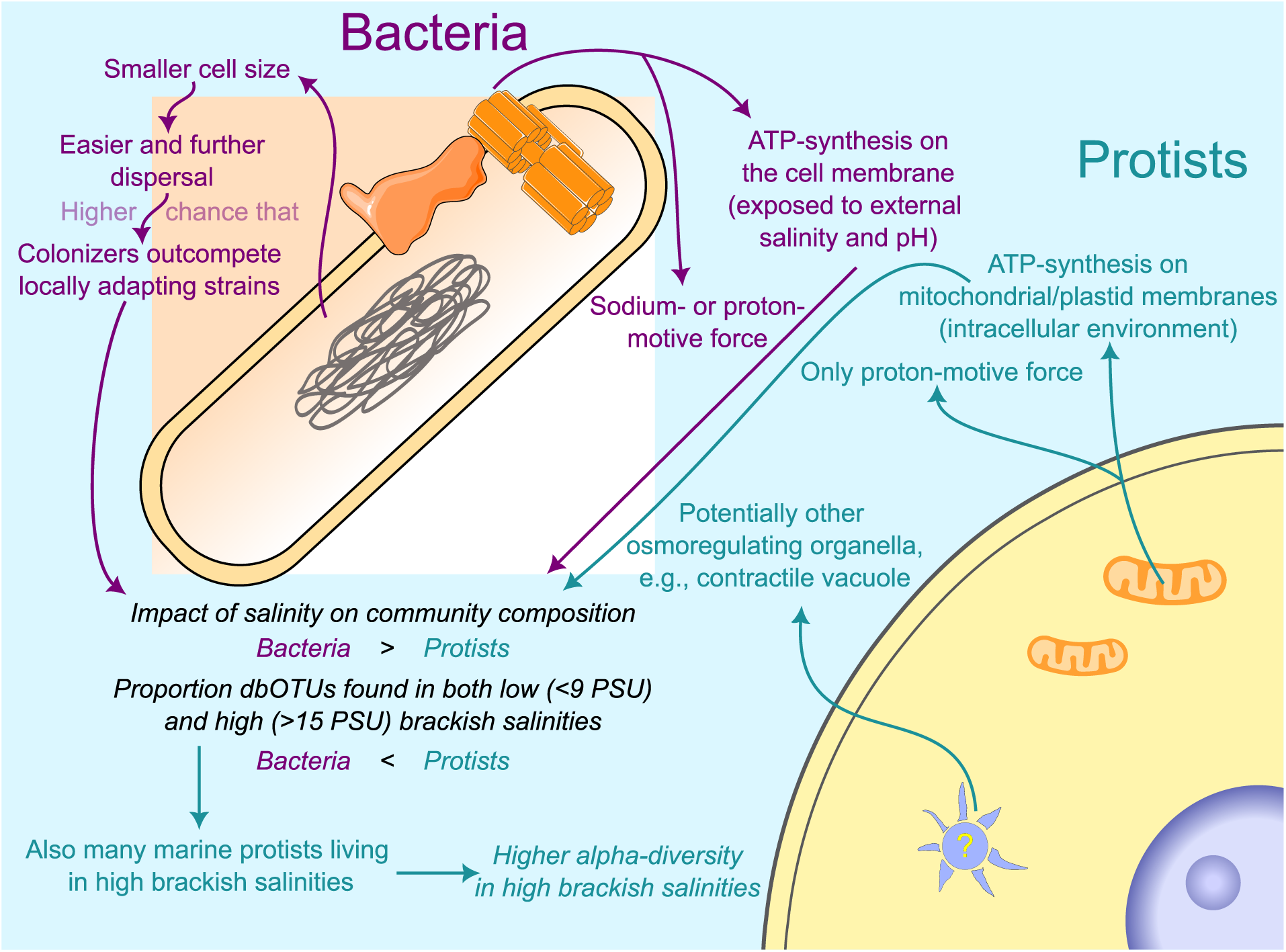
The proposed mechanistic explanation for higher ecological sensitivity to salinity for bacteria than for protists. The cells and their components are drawn schematically and do not accurately represent sizes and shapes. All the cell components, except the protist nucleus and bacterial nucleoid, correspond to the machinery responsible for processes indicated by the arrows leading from them. Statements based on our results are written in italics. This figure was created using images from Togo Picture Gallery^85^ Servier Medical Art (http://smart.servier.com).

The bioenergetics-based explanation would imply that eukaryotes are also ecologically less sensitive to pH. Indeed, that has been observed when comparing bacterial and fungi soil communities^82^. The role of bioenergetics would also imply testable predictions. One is that adding a salt mix that includes (bi-)carbonates, compared to adding pure NaCl, should have a substantially larger impact on bacterial communities, but less so on protist ones. This is expected since (bi-)carbonates introduce a pH buffering and alkalizing effect, neutralizing the protons exported outside the outer membrane. Another prediction would be that the shifts in community composition along the salinity gradients largely follow a shift towards more bacteria using SMF and less using PMF. However, PMF has bioenergetic advantages, and even some halophilic bacteria developed adaptations to use it^83^. Similarly, the bacteria that recently transitioned to higher salinity rarely acquired the bioenergetic machinery to maintain SMF^29^. Thus, SMF-independent adaptations may also allow bacteria to thrive in higher salinities. Nevertheless, around 70% of marine bacteria were predicted to use SMF based on their genomes, compared to 4% in freshwater^27^. The same SMF-producing genes increase in abundance in metagenomes across the Baltic Sea salinity gradient^18^. How well bioenergetic strategy shifts explain the salinity-associated taxonomic turnover is yet to be determined.

In addition to compartmentalization, protist plankton seem to be more constrained in their geographic ranges than bacteria^28,84^. In general, dispersal limitation increases with the size of a planktonic organism^84^. Thus, smaller bacterial cells reach new environments more frequently and from further away. In turn, a locally adapting bacterium will, more likely than a locally adapting protist, be outcompeted by a colonizer from a distant yet environmentally similar location. Previous research shows that the same brackish bacterial species more often inhabit distant brackish basins than closely located freshwater or marine waters^19,29^. This might be less common for protists. Our observations that protists display less community structuring by salinity, more frequently cross the salinity barrier, and exhibit increasing species richness with salinity, are all compatible with local adaptation playing a more important role for protistan than bacterial distribution in the region. More direct, genomic studies are needed to test whether local protist adaptations after a shift of salinity regime are indeed more common than bacterial ones.

An important aspect of testing those hypotheses is that ecological sensitivity does not necessarily equal physiological sensitivity in laboratory conditions. An organism that can survive a wide range of salinities in isolation might lose its competitive edge after a minor disruption.

## Conclusions

We show multiple lines of evidence that salinity, generally, has a stronger impact on the distribution of bacteria than protists. Both patterns of overall community composition and the likelihood of crossing a salinity barrier support this notion. This cannot be generalized to stronger environmental filtering among bacteria, as other environmental factors had similar or stronger effects on the protist communities. To explain these patterns, we speculate that compartmentalization allows protists to acclimatize to a different salinity regime more easily (Fig. 7). Additionally, more frequent and distant dispersal of bacteria may hamper local adaptation, as locally adapting strains are outcompeted by colonizers.

Bacterial alpha diversity followed primarily seasonal patterns. We connected those patterns to changes in water column stratification by detecting deepwater taxa at the surface in vertically mixed winter waters. Moreover, the total relative abundance of deepwater taxa reached substantial levels in winter. While seasonal vertical mixing has previously been connected to higher bacterial alpha diversity^33,44,66,67^, we provide more conclusive evidence of an influx of deepwater taxa to the surface in winter. The impacts of these community coalescence events on surface water ecosystems require further investigation.

In contrast to bacteria, protist alpha diversity primarily followed a geographic pattern, being higher in the more saline Kattegat-Skagerrak. This pattern is analogous to that of metazooplankton and fish, which originates from the limited scope of marine taxa’s adaptations to the Baltic Sea conditions following the last glaciation period^11,20^. Whether the diverse protists in the Kattegat-Skagerrak represent marine taxa that did not adapt to lower salinities in the Baltic Sea is yet to be confirmed.

We show geographic differences not only in microbial community composition but also in seasonal succession dynamics. Thus, time series results from multiple and diverse locations should be integrated for generalizability and proper interpretation. We observed interannual stability of community composition relative to the environmental conditions, consistent with previous results. However, minor technical differences between the datasets confounded alpha diversity analysis. Thus, a standardized DNA-based scheme is needed to monitor rare microbial taxa.

Using DNA metabarcoding across strong spatiotemporal gradients, we find new patterns of microbial diversity and expand our understanding of previously observed. Ultimately, as Earth’s environments undergo rapid change, diverse microorganisms, often understudied groups, will be impacted and/or consequential for the trajectory of coastal and other ecosystems. DNA-based monitoring covers the phylogenetic scope needed to understand and respond to those unprecedented changes.

## Methods

### Data collection and sequencing

The 2019-2020 data comes from Latz et al. 2024^45^. The collection and sequencing procedures are described in detail therein. In brief, samples were collected monthly or every second week, depending on the location, as a part of the Swedish National Marine Monitoring Program. The samples for metabarcoding were collected using a depth-integrating hose, covering 0-10m (except 0-5m at RÅNEÅ-1, and 0-20m at B1 and BY31). 500 ml of water from the hose sampling was filtered using 0.22 µm pore size. DNA was extracted using the Zymobiomics™ DNA miniprep kit following a modified protocol^86^, and spike-in DNA was added. The DNA was sequenced using Illumina™ MiSeq flow cells. For 16S rDNA gene-based metabarcoding of prokaryotes, primers targeting the hypervariable V3-V4 regions^16^, and for 18S metabarcoding of eukaryotes, primers targeting the V4 region^87^, were used. Phased primers^45,88^ were used for both 18S (on the forward primer) and 16S (on both primers) metabarcoding. Sequences of the primers and the details of the phasing strategy and sample indexing through a second PCR can be found in Latz et al. 2024^45^. Physico-chemical and geographic data has been obtained from the Swedish Meteorology and Hydrology Insitute’s service SHARKweb: https://sharkweb.smhi.se/hamta-data/

The 2015-2017 data was used for the first time for this study. The samples were stored at -80°C. DNA was extracted in the summer of 2017 and stored at -20°C until February 2023. The sampling and sequencing procedures were analogous to the one for 2019-2020 data, with a few exceptions:

1. Volumes varied from 200ml to 800ml. According to our previous tests run in the same setting, volumes ≥200ml should capture the microbial community structure equally well^45^.
2. DNA extraction was performed using Qiagen™ DNeasy PowerWater Kit, and not ZymoBIOMICS™ DNA Miniprep Kit with a small modification^86^ as for the 2019-2020 dataset.
3. No spike-in DNA was added.
4. Illumina™ NextSeq, rather than MiSeq system, was used.
5. For 18S, the initial use of 20 cycles for the first PCR amplification was insufficient, leading to many library preparation failures. It was then re-run with 24 cycles, and the results of both runs were merged after processing the tables by summing up the counts in the output count table.
6. For 18S, reverse primers were also phased, following a sequence (CTACGA)CTTTCGTTCTTGATYRR, with versions of the primer starting from any base in the brackets included in the primer mix.
7. After library preparation, for most samples, 18S and 16S libraries were mixed and sequenced together. A cutadapt^89^-based pipeline was used to separate reads with only 16S or only 18S samples for downstream analysis, as described in the following section.

Samples from other projects, which we did not analyze in this study, were sequenced simultaneously and submitted to the European Nucleotide Archive (ENA) all together.

### Processing of sequencing data

The data processing for 2019-2020 samples has been described in detail in Latz et al. 2024^45^. In brief, the phased primer sequences were removed from the obtained reads using a modified version of a cutadapt^89^-based pipeline (original pipeline taken from: https://github.com/kjurdzinski/amplicon-multi-cutadapt-18S-16S-monitoring). Only reads containing 16S or 18S primers were kept, which allowed us to separate reads from the two methods when the libraries were mixed. DADA2^90^ version 1.18.0 was used to denoise the data, infer amplicon sequence variants (ASVs), and taxonomically annotate them. For taxonomic annotation of 16S ASVs, we used 16S sequences from GTDB^91^ version R06-RS202-1 corrected for mislabeled sequences using SATIVA^92,93^. Annotation was additionally conducted using the SILVA database^94^ version 138.1 to identify plastids and mitochondria. The 18S ASVs were annotated using the PR2 database^95^ version 5.0.1.

The 2015-2017 data was analyzed the same way, with the only difference being that the denoising and ASV inference with DADA2 was performed using the nf-core/ampliseq pipeline^96^ version 2.7.0 instead of an in-house R script. We kept the DADA2 parameters as in Latz et al. 2024^45^.

We further combined the 2015-2017 and 2019-2020 data. We matched ASVs using exact matching. On the merged ASV-sequence list and count table, we ran an additional chimera removal step using UCHIME v1^97^ and clustered the remaining ASVs based on their simultaneous genetic and distribution similarity using dbOTU3^46^. Both these steps were performed using an in-house pipeline available at https://github.com/kjurdzinski/chimera_dbOTU_pipeline. Default parameters were used for UCHIME, and for dbOTU2, only the maximum genetic distance for 16S has been changed to 3% (instead of the default 10%). The procedure should minimize the influence of intra-specific variation on our data and further reduce the effects of PCR amplification and sequencing errors. We called thus obtained taxonomic units dbOTUs (distance-based operational taxonomic units, as inferred using dbOTU3^46^)

For both 16S and 18S, we removed dbOTUs with no annotation at the phylum/supergroup taxonomic level. From 16S dbOTUs, *Archaea*, plastids, and mitochondria were removed. Thus, the remaining ASVs should correspond to bacteria. From 18S ASVs, animals (*Metazoa*), *Fungi*, land plants (*Embryophyceae*), and selected macroalgae (*Phaeophyceae*, and Rhodophyta excluding *Rhodella* and *Cyanidiales*) were removed. Thus, the remaining ASVs were primarily protists.

We kept spike-in dbOTUs in a separate table and used them only to obtain spike-in corrected abundance, i.e., the inferred number of rDNA copies per liter. The rDNA copies per liter were calculated according to the equation:

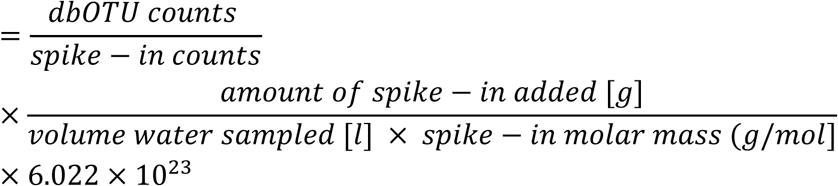

We have excluded over- and under-sequenced samples to remove the amplification errors (including distorted relative abundances) and not to rarefy to a too low read number during rarefaction. We have excluded three samples with less than 30,000 reads for bacteria, as well as one sample with more than 2,000,000 reads. Moreover, we have excluded two samples from 2019-2020 with more than 200,000 protist reads (this number of reads was normal for 2015-2017 data since Illumina™ NextSeq was used instead of MiSeq, yielding overall higher numbers of reads). Additionally, one sample (20191015_263953_1) with suspected contamination has been excluded.

For comparability of results for protists and bacteria, we used only the samples with data available from both 16S and 18S amplicon sequencing (excluding the under- and over-sequenced samples) for all the further analyses. Additionally, in two samples (station RÅNEÅ-2 on 2019-01-29 (sample ID: 20190129_263856_1) and B7 on 2019-09-17 (sample ID: 20190917_264450_1)) 18S spike-in addition failed. Those samples were excluded from analyses in which spike-in normalization was used, except for bacteria-only focused analyses in Fig. 1b.

### Abundance, alpha and beta diversity spatiotemporal patterns

For the spatiotemporal analysis of abundance, beta- and alpha diversity across the samples from 2019-2020, we removed the samples from stations H4, BY29 / LL 19, and SR3. Those stations were sampled at only a few unevenly distributed occasions, and their exclusion/inclusion at certain seasons could obscure seasonal patterns.

All the statistical analyses and data visualization was performed using R version 4.2.2^98^. Inkscape^99^ was used to assemble and edit multi-panel figures, and to make Fig. 7.

The multivariate analyses were performed using vegan R package^100^, and all the functions in this paragraph can be found there unless another package is explicitly stated. Bray–Curtis distances between communities were obtained using rarefaction via the avgdist function. Those distances were used for Principal Coordinate Analysis (PCoA) and distance-based Redundancy Analysis (dbRDA), performed with, respectively, capscale (without setting constraints) and dbrda functions. Variance partitioning was performed using rdacca.hp R package^101^. Families were positioned on the ordination plots by calculating weighted averages (sppscores function). Only the families deviating (Bonferroni corrected P-value < 0.05) from normal distribution around the center of the dbRDA coordinate system were shown on the plots. To find the corresponding two-dimensional ellipsoid confidence intervals, aspace^102^ package was used. Whenever only samples were visualized, their ordination scores were scaled using the eigenvalues, and symmetric scaling was used if both the families and the samples were visualized (using, respectively, “sites” and “symmetric” scaling options of the plotting functions respectively). For dbRDA and variance partitioning, we used a subset of 228 (out of 246) samples with all the environmental variables of interest available. For more informative visualization, to keep the sample structuring visible despite very high family scores, we divided the family scores in half.

dbOTU richness was calculated as the mean number of dbOTUs detected in a sample over 100 rarefaction iterations. Variance partitioning was performed using rdacca.hp R package^101^.

For both beta and alpha diversity estimation, in each rarefaction iteration, we rarefied the bacterial 16S reads to 34382 per sample, and the protists 18S reads to 8084 per sample.

For deepwater taxa detection, we used the representative sequences of 99% average nucleotide identity OTUs (operational taxonomic units) detected in samples from the summer of 2008 and coming from a previous study^33^. The 2008 summer transect included sampling across different depths, and the deepwater OTUs in this study were present exclusively at depths ranging from 11 to 300m, classified as mesopelagic in the original study^33^. We connected the OTUs from the original, transect-based study to ASVs from the monitoring data used here, using the 99% nucleotide identity threshold in VSEARCH^68^. We deemed an OTU and a dbOTU to be matching if their representative sequences (most abundant ASV for a dbOTU) matched in the VSEARCH output. For comparing the number of OTUs from deepwater, surface water, and both (Fig. 5a) in winter and summer we used the chi-squared test as implemented in the prop.test function in R^103^.

### Proportion salinity barrier crossing dbOTUs and Bayesian modelling

Any dbOTU found among included samples with salinity >15 PSU and among samples with salinity <9 PSU was regarded as barrier crossing. For balancing out the number of samples on both sides of the salinity barrier, apart from the four stations with salinity >15 PSU (Å17, SLÄGGÖ), four out of twelve stations with salinity <9 PSU: BY2 ARKONA, BY5 BORNHOLMSDJ, BY15 GOTLANDSDJ, REF M1V1 (see Fig. 1a for locations of the stations). First, we chose the four most saline stations from among those with salinity <9 PSU. Then, we switched BCS III-10 to REF M1V1 (see Fig. 1), since the latter had more samples, and, unlike other chosen stations from lower salinities, was located close to the mainland coastline. For each station but the least sampled (REF M1V1), the samples were downsampled, choosing samples closest in time to the ones collected at the least sampled stations. Whenever available, those samples should correspond to the same monitoring cruises.

The Bayesian models were built and implemented in JAGS ^74^, using the R package rjagS10^104^. The parameters, probabilities, and predictions were calculated based on every fifth out of 10000 MCMC iterations.

The model for probability of crossing the salinity barrier was optimized based on Bernoulli distribution:

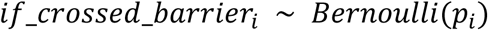

where for each dbOTU *i*, *if*_*crossed*_*barrier*_*i*_ is a binary variable = 1 if it is a barrier crossing dbOTU. *p*_*i*_ was calculated according to the formula

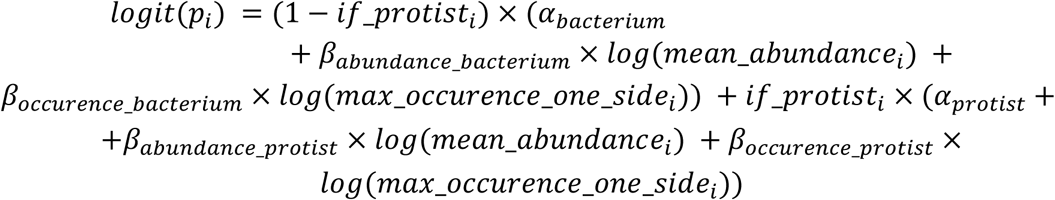

where *if*_*protist*_*i*_ is a binary variable = 1 if the dbOTU *i* is a protist. *mean*_*abundance*_*i*_ is geometric mean of the abundance of the dbOTU *i* in the samples where it was present. *max*_*occurence*_*one*_*side*_*i*_ is the maximum out of the numbers of samples in which the dbOTU *i* was found at one side of the salinity barrier (in salinity <9 PSU or >15 PSU). The other variables are the parameters of the model.

We compared this model with one assuming the same abundance and occupancy dependence for bacteria and for protists (one *β*_*abundance*_ and *β*_*occupancy*_, same for protists and bacteria, different *α*_*bacterium*_and *α*_*protist*_). For both the model with different and the same abundance and occupancy dependencies, we also compared the versions without log-transforming either only the occupancy, or both abundance and occupancy values. Based on the Watanabe-Akaike criterion^75^, the model we used performed the best.

The model for total occupancy was build based on the Poisson distribution

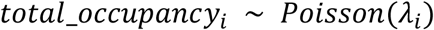

where *total*_*occupancy*_*i*_ is the total number of samples in which dbOTU *i* was found. *λ*_*i*_ was calculated according to the formula:

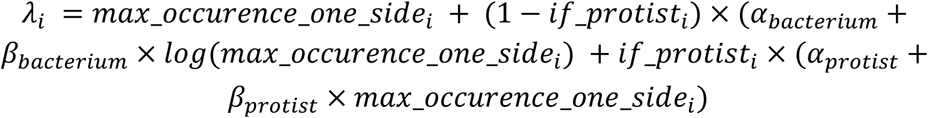

with *if*_*protist*_*i*_ and *max*_*occurence*_*one*_*side*_*i*_ having the same meaning as described above. The occupancy at the less occupied side of the salinity barrier was calculated by subtracting *max*_*occurence*_*one*_*side* from the predicted *λ*.

## Acknowledgements

We would like to thank Jenny Lycken for the effort put into sampling and sample preparation for the 2019-2020 dataset. National Genomics Infrastructure (NGI) Sweden, and especially Elísabet Einarsdóttir, for seamless and adjustable sequencing service. Karin Garefeld for help with preparing the 2015-2017 samples for sequencing. Additionally, KTJ would like to thank Maliheh Mehrshad for hosting him in Uppsala during the critical period of work on this manuscript, as well as suggesting courses which taught him the skills necessary for the crucial analyses of this study. Ulf Gardin, for help with the multivariate analyses. Mathew Low and Malin Aronsson for help with Bayesian modeling.

## Study funding information

This work was funded by the Swedish Agency for Marine and Water Management and the Swedish Environmental Protection Agency under grant number NV-03728-17. AFA was additionally funded by a research grant (2021-05563) from the Swedish Research Council VR. DNA-extraction and sequencing of the 2015-2017 samples was supported by the EU Horizon 2020 project JERICO-NEXT under grant agreement No 654410 and by the Swedish Research Council FORMAS (grant number 214-2013-1449).

For this study, we used computational resources provided by the Swedish National Infrastructure for Computing (SNIC) through the Uppsala Multidisciplinary Center for Advanced Computational Science (UPPMAX).

## Author contributions

KTJ analyzed the data under the supervision of AFA and with help from MACL and AT. YOOH prepared the DNA for the 2015-2017 dataset. KTJ wrote the first version of the manuscript under supervision of AFA. All authors contributed to conceptualization, interpretation of the data, revised the manuscript, and approved the final version.

## Data and code availability

The 2019-2020 data have been published before as a data paper^45^. The 2015-2017 raw sequencing data is available in the European Nucleotide Archive under the study accession code https://identifiers.org/ena.embl:PRJEB84926. The ASVs identified in the 2015-2019 sequencing data, together with a count table and contextual data, can be accessed through the ASV portal at the Swedish Biodiversity Data Infrastructure (SBDI). Subsequently, the ASVs, count tables, and contextual information are available at the Global Biodiversity Information Facility (GBIF). The code used to analyze the data and generate the figures, together with intermediate processing output, can be found at https://doi.org/_/scilifelab.

## Supplementary figures

**Supplementary Fig. S1.**
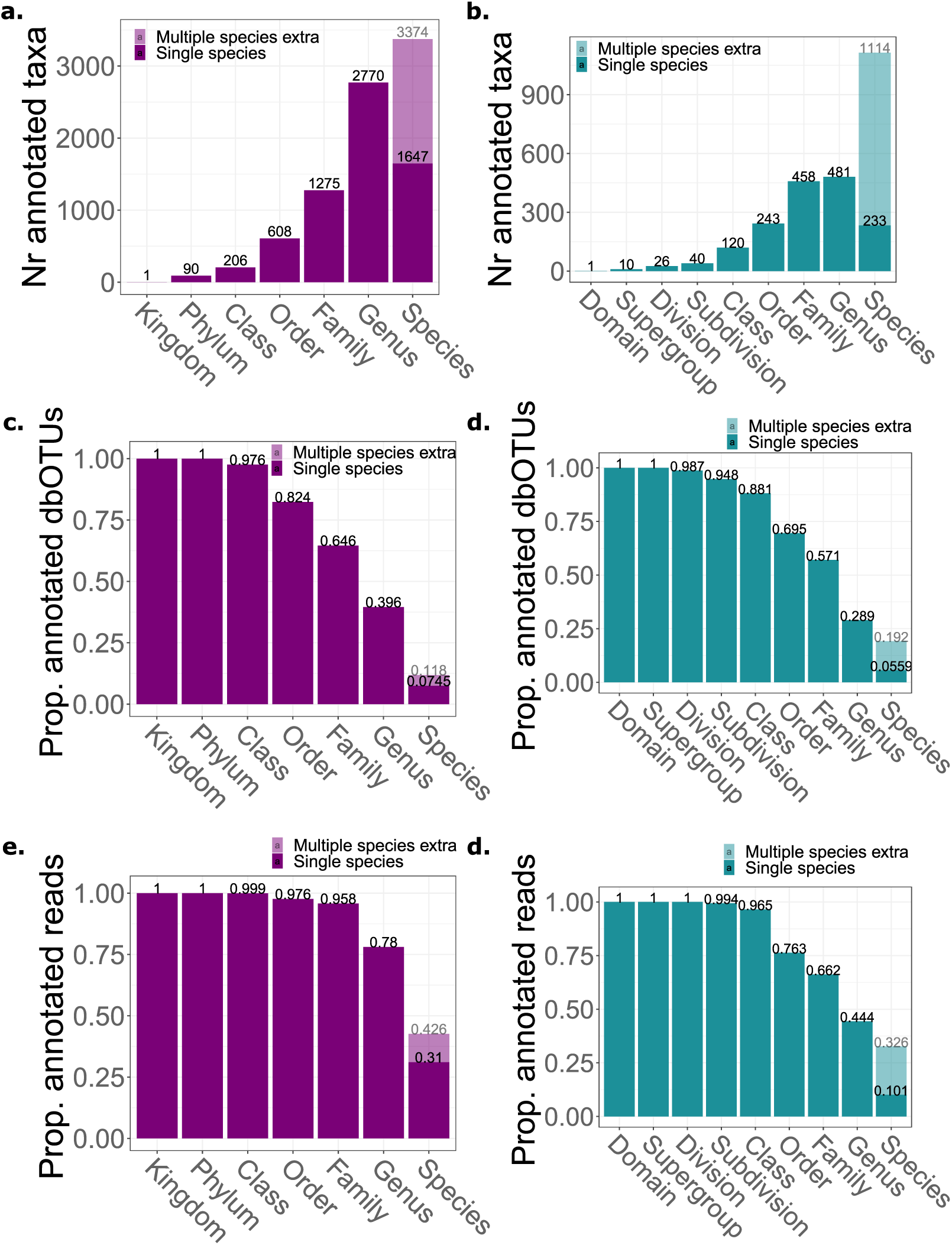
Taxonomic annotation statistics. **a-b**. Number of different bacterial (**a.**) and protist (**b.**) taxa found at different taxonomic levels. **c-d**. Proportions of bacterial (**c.**) and protist (**d.**) taxa annotated at different taxonomic levels. **e-f**. Proportions of bacterial (**e.**) and protist (**f.**) reads annotated at different taxonomic levels. “Single species” corresponds to an analysis that provides a species-level annotation only if the query sequence has identical matches to database sequences of a single species. “Multiple species extra” is the extra number/proportion of annotated species when exact matches to multiple species are allowed.

**Supplementary Fig. S2.**
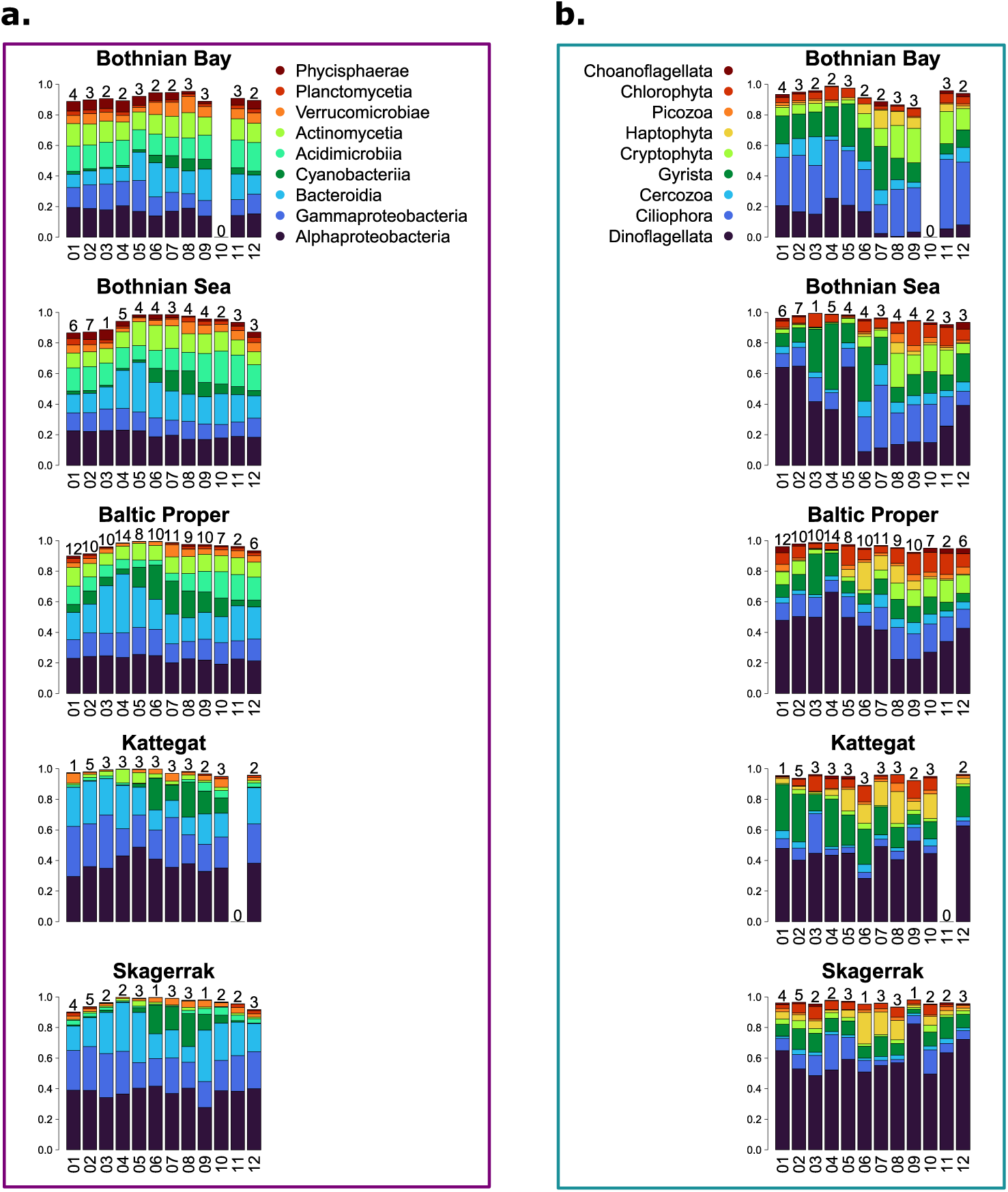
Spatiotemporal changes in abundance of major microbial taxa, 2019-2020. **a-b.** The proportion of rDNA marker gene reads annotated to **a.** bacterial classes and **b.** protist subdivisions averaged over each month and Baltic Sea area basin. Only the classes/subdivisions which, on average, corresponded to >0.01 of reads across samples are shown. Above each bar, the number of samples collected in the respective month and basin is given.

**Supplementary Fig. S3.**
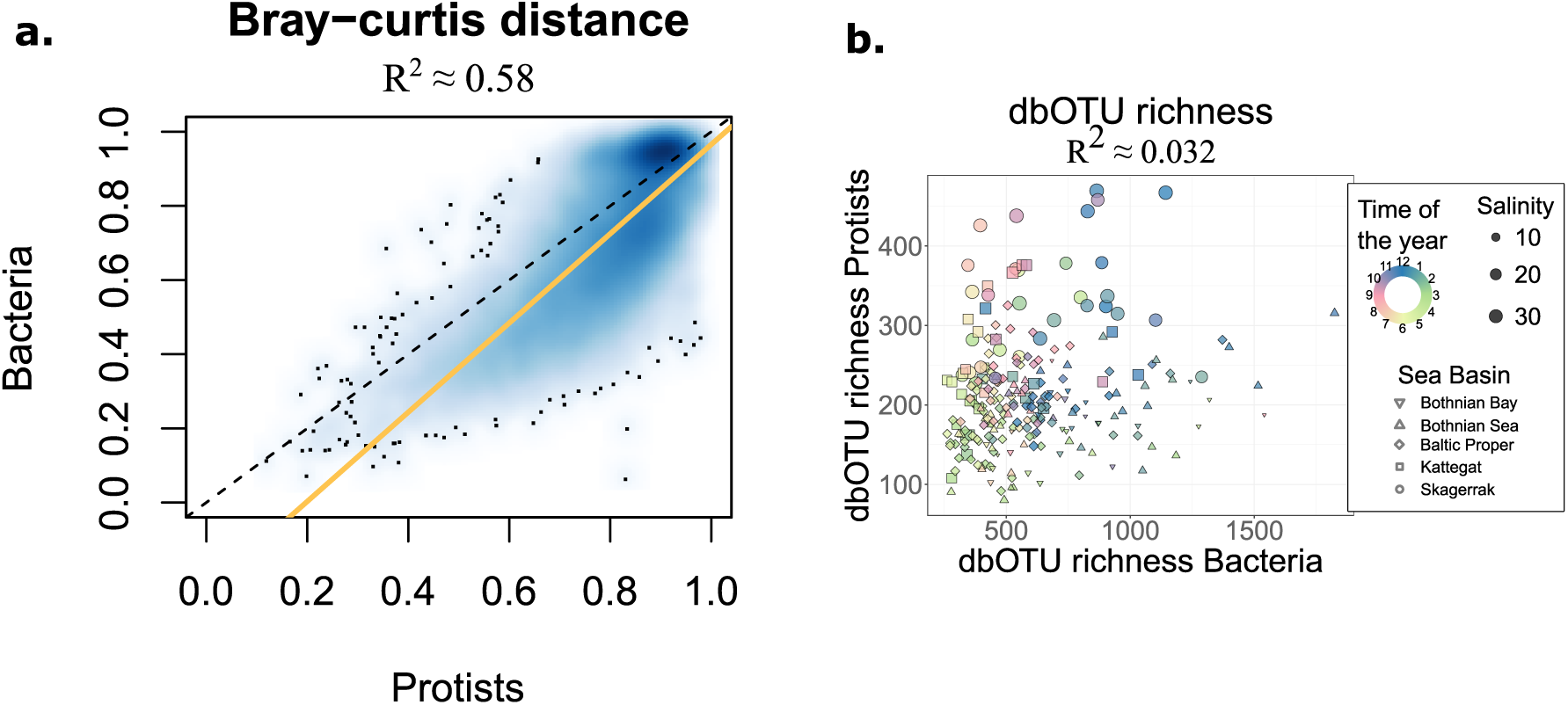
Dissimilarity of diversity metrics between bacteria and protist, 2019-2020. **a.** Beta diversity calculated as Bray-Curtis distances. The golden line represents the best linear fit, while the dashed black line is the identity line. **b.** Number of detected dbOTUs (alpha diversity).

**Supplementary Fig. S4.**
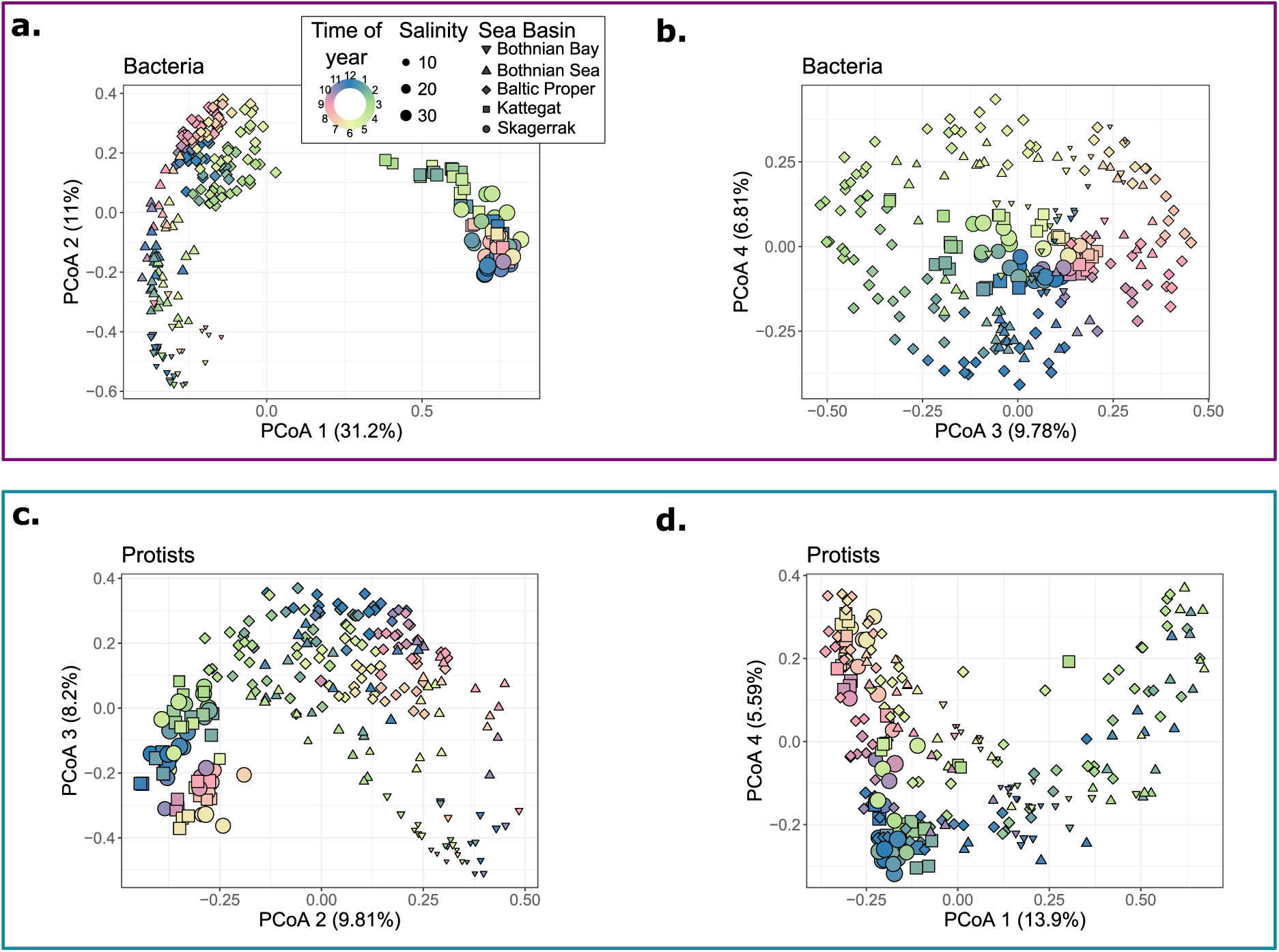
Principal coordinate analysis (PCoA) 2019-2020. The first four ordination axes are based on Bray-Curtis distances between bacterial (**a-b**) and protist (**c-d**) communities. Note: The axes are grouped and ordered by which ones, after visual examination, correspond to geographic structuring (**a**, **c)** and which ones to seasonal effects (**b**, **d**). This ordering corresponded to showing ordination axes with decreasing variation explained for bacteria (**a-b**). However, for protists, the first plot (**c**) shows the second and the third components, while the second plot (**d**) shows the first and the fourth components.

**Supplementary Fig. S5.**
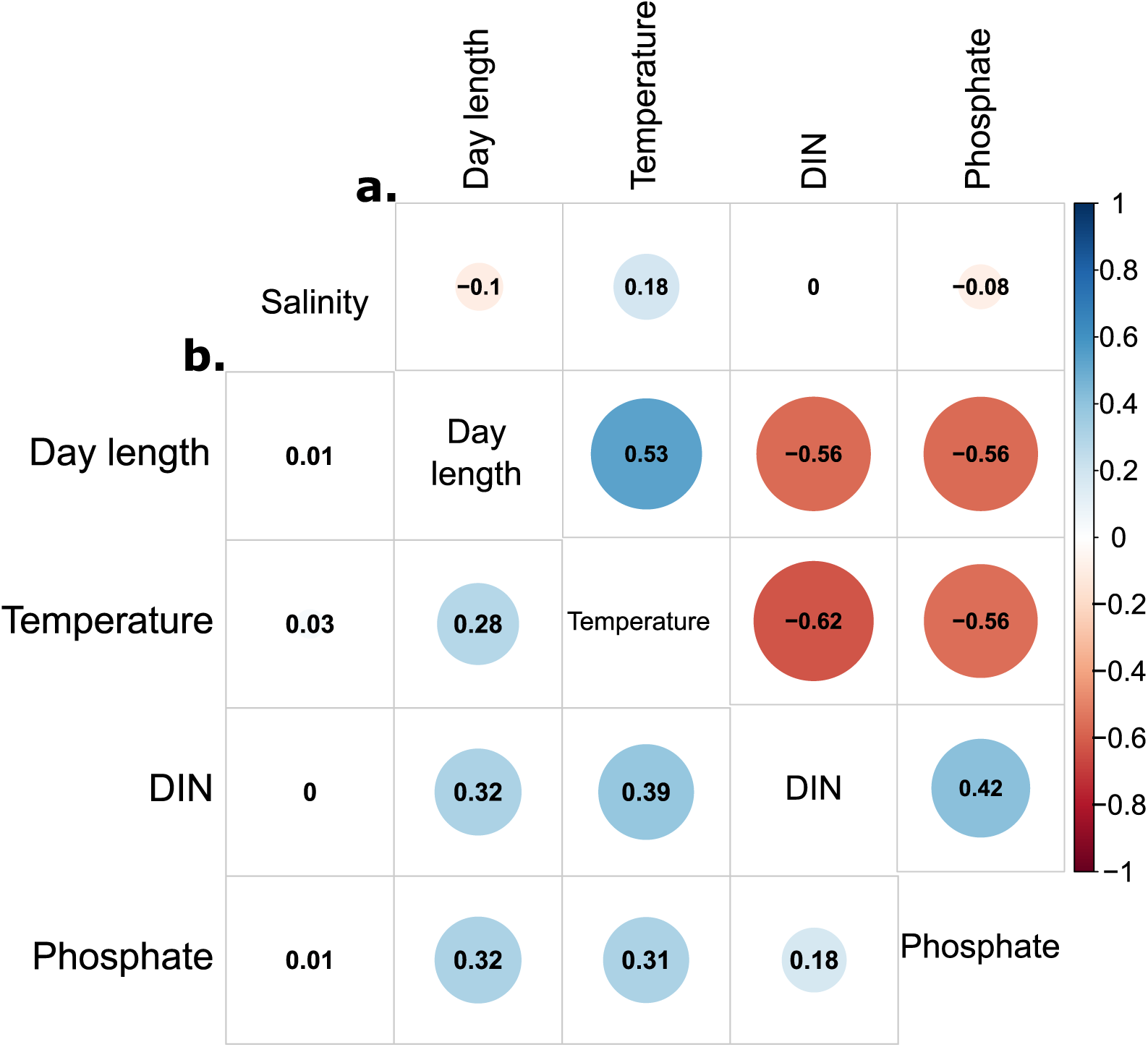
Correlations between chosen environmental variables for the 2019-2020 dataset. **a.** Pearson correlation coefficients (ρ) on an upper-triangle correlation plot. **b.** Coefficients of determination (R^2^) on a lower-triangle correlation plot.

**Supplementary Fig. S6.**
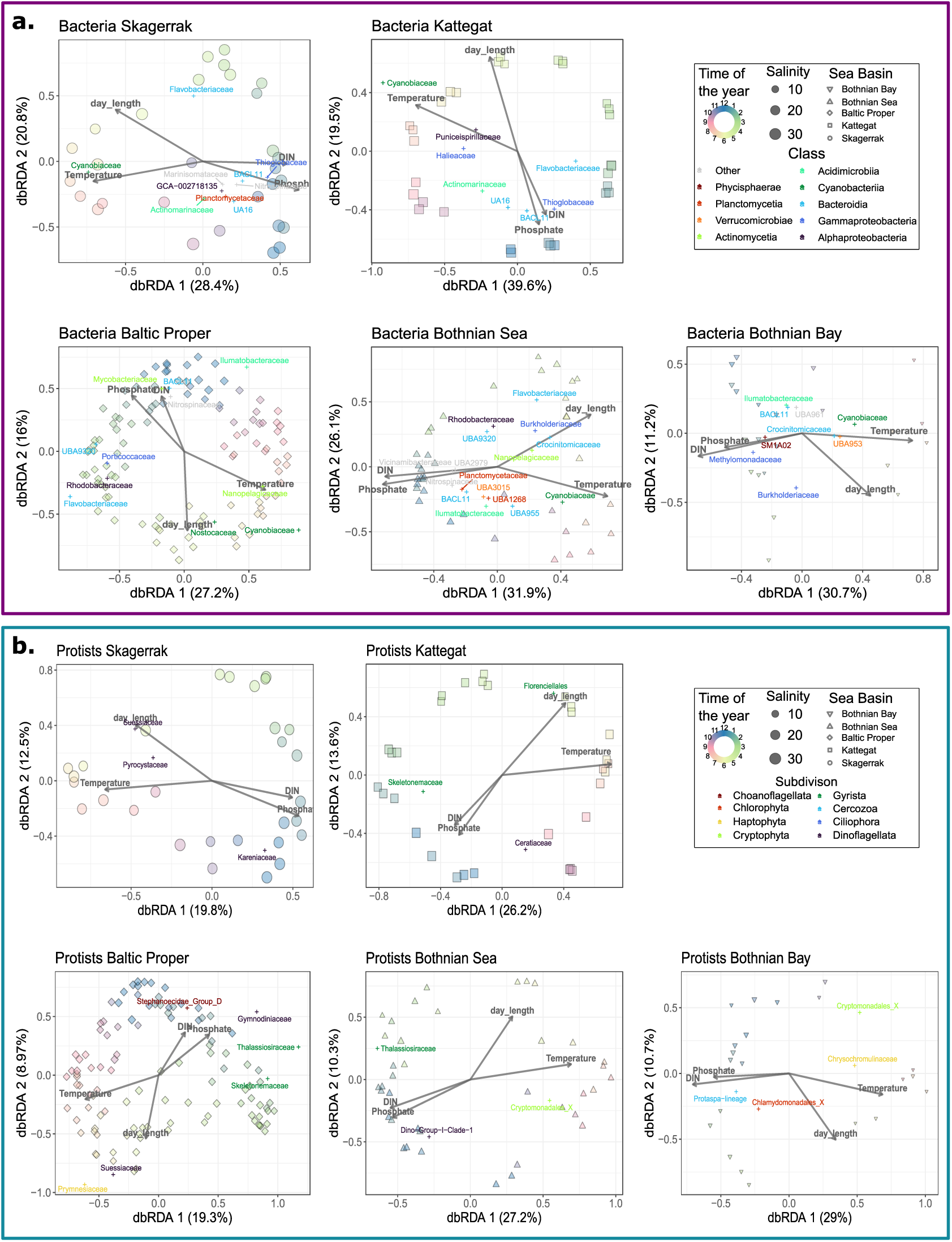
Seasonal changes in community composition within distinguished basins of the Baltic Sea area. distance-based Redundancy Analysis (dbRDA) based on Bray-Curtis distances between the bacterial (**a.**) and protist (**b.**) communities within each of the distinguished basins of the Baltic Sea area. Weighted averages of families deviating (Bonferroni corrected P < 0.05) were placed on the plot, with values rescaled by a factor of 0.5. Percentages of variation explained by each dbRDA component are given in brackets by the axes labels.

**Supplementary Fig. S7.**
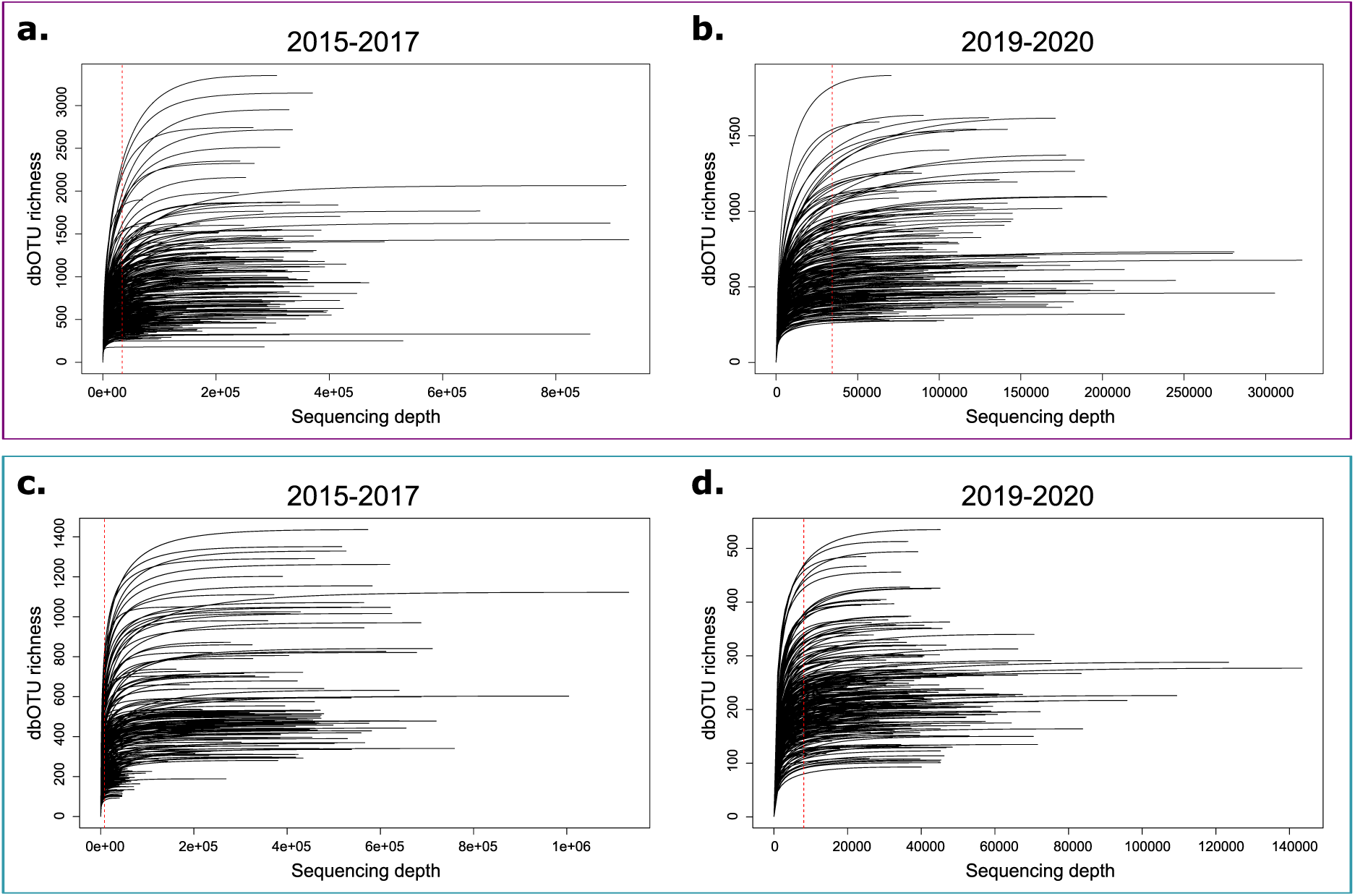
Refraction curves. **a-b**. Based on 16S filtered (only bacteria) read from the 2015-2017 (**a.**) and 2019-2020 (**b.**) datests. **c-d**. Based on 18S filtered (only protists) read from the 2015-2017 (**c.**) and 2019-2020 (**d.**) datests. dbOTU - distribution-based operational taxonomic unit.

**Supplementary Fig S8.**
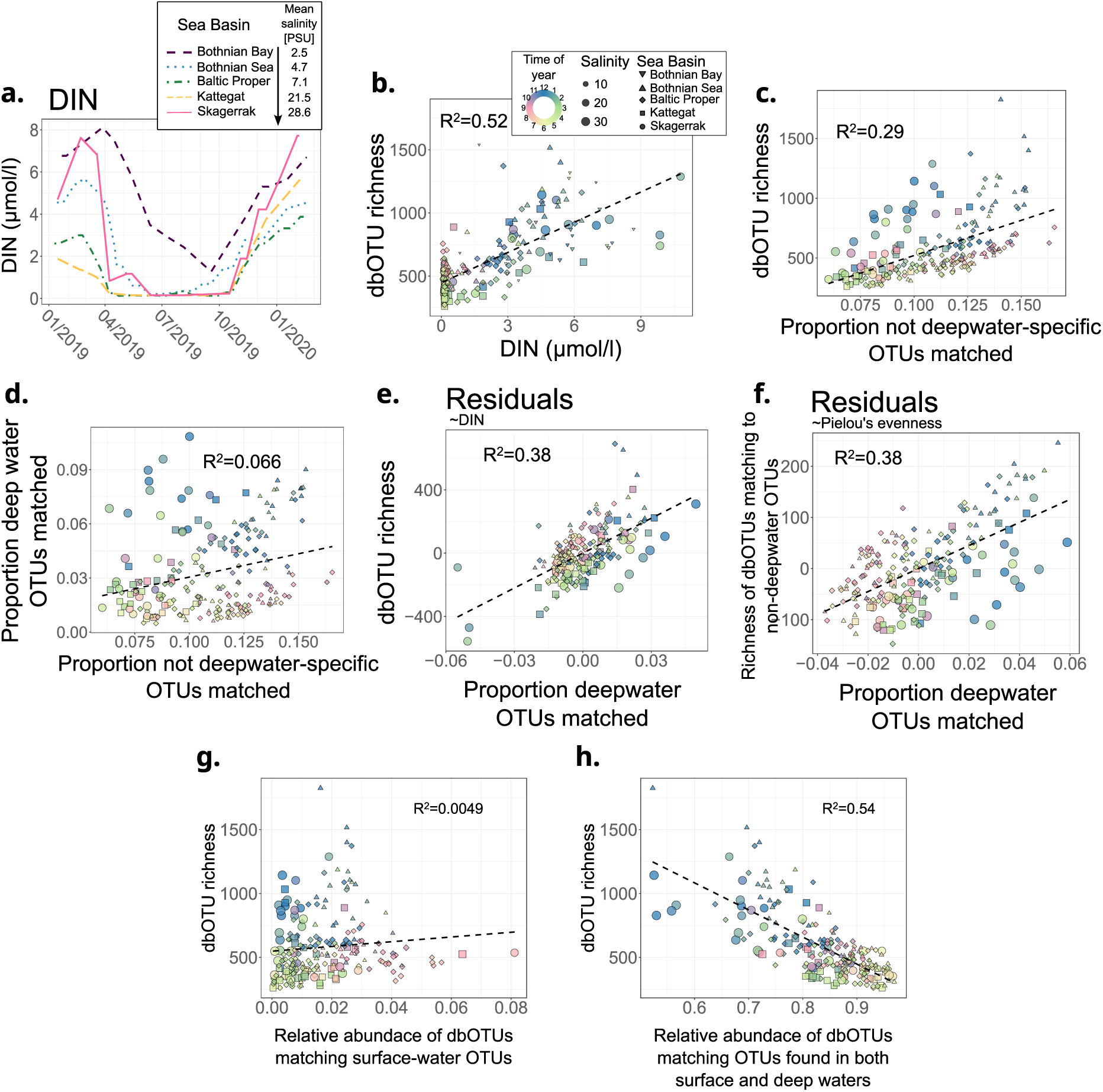
Bacterial alpha diversity. **a.** A rolling mean of DIN values, averaged for measurements two weeks before and after each date. **b**. DIN vs. dbOTU richness across samples. **c**. Proportion of all non-deepwater-specific OTUs (“surface” and “both” categories) having a match to a dbOTU from a sample vs. bacterial dbOTU richness. **c**. Proportion of all non-deepwater-specific OTUs vs. proportion of all deepwater OTUs. **e.** The residuals of proportion of all deepwater OTUs vs. dbOTU richness from models correlating the factors to DIN, corresponding to partitioning out the unvisualized parameter. **f. e.** The residuals of proportion of all deepwater OTUs vs. dbOTU richness from models correlating the factors to Pielou’s evenness. **g.** Relative abundance of all dbOTUs matching a surface water OTU vs bacterial dbOTU richness. **e.** Relative abundance of all dbOTUs matching an OTU found in both surface and deep waters vs bacterial dbOTU richness. Bothnian Bay was excluded from plots c-g, since there were no deepwater samples taken in this region in the transect-based study^33^. The same legend applies to figures b-g. DIN - dissolved inorganic nitrogen.

**Supplementary Fig. S9.**
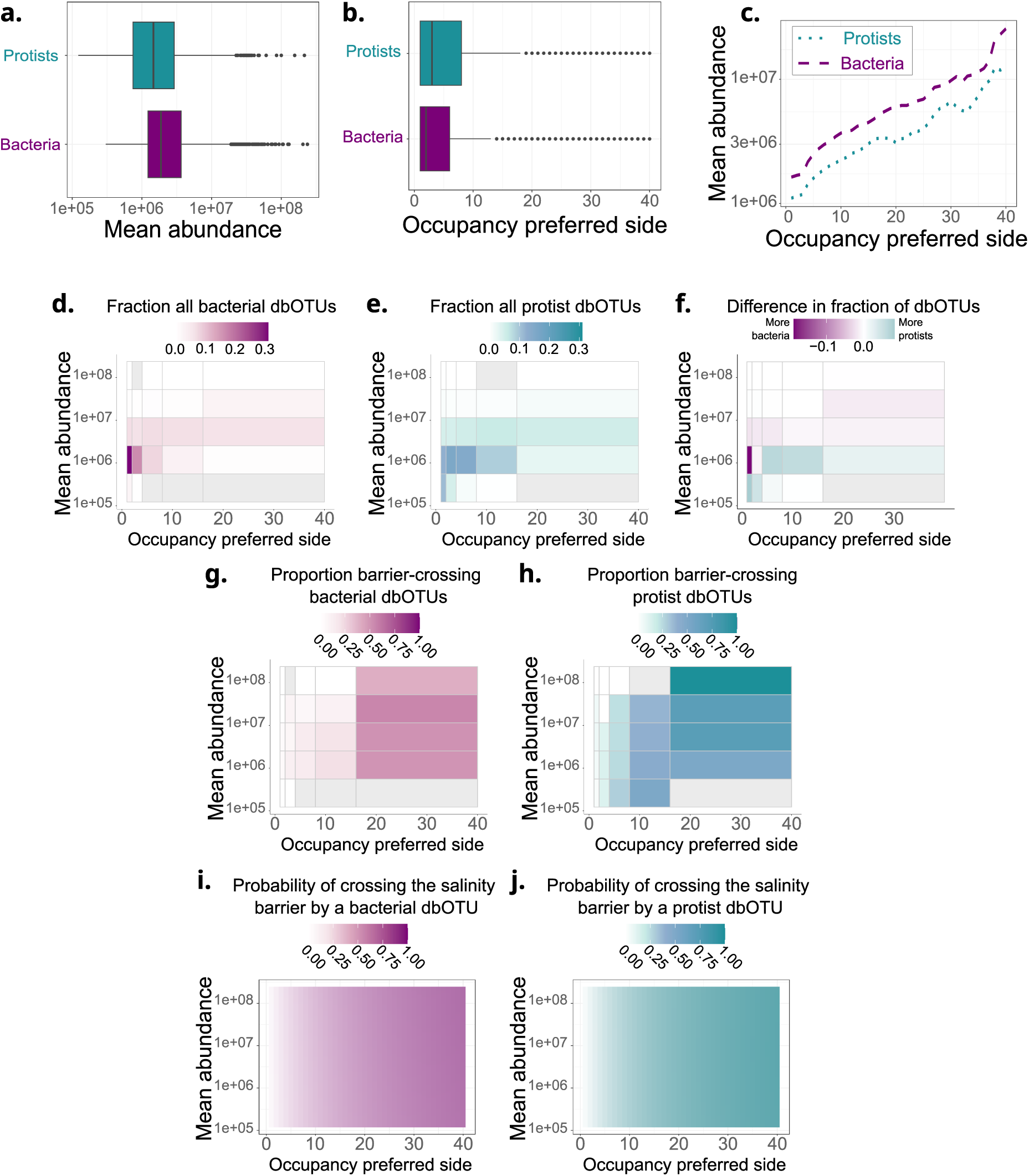
The fractions of all dbOTUs, the proportions of barrier-crossing dbOTUs, and the probability of crossing the barrier across occupancy and abundance values. **a-b.** The distribution of mean abundance (**a.**) and occupancy at the preferred side of the salinity barrier (the side where the dbOTU has higher occupancy) (**b.**). **c.** Rolling mean (+/- two bins) of abundance across the occupancy values. **d-f.** The fraction of all bacterial (**d.**) and protist (**e.**) dbOTUs, and the difference between the fraction of protist and bacterial dbOTUs (**f.**). **g-h**. The proportion of barrier crossing bacterial (**g.**) and protist (**h.**) dbOTUs. **d-h.** Grey tiles correspond to the values across which either no protist or no bacterial dbOTUs were found. **I-j.** The predicted probabilities of a bacterial (**i.**) and protist (**j.**) dbOTU crossing the salinity barrier. For all the analyses, only the four stations with salinity >15 PSU, and four out of the most saline stations with salinity <9 PSU are chosen. The data for each station is downsampled to the same number of reads. During downsampling, observations closest in time to the samples from the least sampled station were chosen. Only observations from 2019-2020 are included. The abundance values correspond to the numbers of rDNA copies per liter, based on the spike-in normalization, and are always given on a logarithmic scale. “Occupancy preferred side” is the maximum occupancy at one of the two sides of the salinity barrier.

